# 2D and 3D multiplexed subcellular profiling of nuclear instability in human cancer

**DOI:** 10.1101/2023.11.07.566063

**Authors:** Shannon Coy, Brian Cheng, Jong Suk Lee, Rumana Rashid, Lindsay Browning, Yilin Xu, Sankha S. Chakrabarty, Clarence Yapp, Sabrina Chan, Juliann B. Tefft, Emily Scott, Alexander Spektor, Keith L. Ligon, Gregory J. Baker, David Pellman, Peter K. Sorger, Sandro Santagata

## Abstract

Nuclear atypia, including altered nuclear size, contour, and chromatin organization, is ubiquitous in cancer cells. Atypical primary nuclei and micronuclei can rupture during interphase; however, the frequency, causes, and consequences of nuclear rupture are unknown in most cancers. We demonstrate that nuclear envelope rupture is surprisingly common in many human cancers, particularly glioblastoma. Using highly-multiplexed 2D and super-resolution 3D-imaging of glioblastoma tissues and patient-derived xenografts and cells, we link primary nuclear rupture with reduced lamin A/C and micronuclear rupture with reduced lamin B1. Moreover, ruptured glioblastoma cells activate cGAS-STING-signaling involved in innate immunity. We observe that local patterning of cell states influences tumor spatial organization and is linked to both lamin expression and rupture frequency, with neural-progenitor-cell-like states exhibiting the lowest lamin A/C levels and greatest susceptibility to primary nuclear rupture. Our study reveals that nuclear instability is a core feature of cancer, and links nuclear integrity, cell state, and immune signaling.

## Introduction

Relative to normal cells, cancer cells exhibit profound alterations in nuclear structure, including irregular shape, size, and multi-nucleation. Cancer cells may also contain aberrant structures such as *micronuclei* and *chromosomal bridges*.^1^ These phenotypes, collectively termed *nuclear atypia*, are among the earliest recognizable features of cancer. Detection of nuclear atypia has been the primary basis for the pathologic diagnosis of dysplasia and neoplasia since the first microscopic descriptions of cancer over 150 years ago.^2,3^

The potential importance of nuclear atypia is underscored by data showing that mis-segregation of chromosomes into micronuclei (MN) and chromosomal bridges underlies the development of chromothripsis, a process of catastrophic chromosomal rearrangements that can drive rapid genomic evolution in many cancers.^4–9^ Chromosomes in micronuclei may additionally accumulate DNA damage and epigenetic alterations through mechanisms that include aberrant base-excision repair.^10,11^ Moreover, micronuclei and chromosomal bridges can suffer rupture of the nuclear envelope (NE), exposing DNA to cytoplasmic nucleases such as TREX1^12,13^ and driving activation of innate immune pathways such as cGAS-STING. This process may contribute to tumor progression by promoting chronic inflammation and conditioning of the tumor immune microenvironment.^14,15^

*Primary nuclei* may also suffer spontaneous or mechanically-induced NE rupture, resulting in uncontrolled exchange of material between the nucleus and cytoplasm.^16–18^ This process may similarly lead to DNA damage and cGAS-STING activation in some settings.^19,20^ Primary nuclear (PN) ruptures are rapidly detected by barrier-to-autointegration-factor (BAF; *BANF1)* and repaired by the ESCRT-III complex.^21^ BAF is a ubiquitously-expressed double-stranded DNA binding-protein essential for nuclear lamina organization and cytoplasmic DNA responses, and BAF localization may serve as a biomarker of NE rupture.^22^ However, despite the strong association of nuclear atypia with cancer, the prevalence of atypical nuclear structures such as micronuclei, chromosomal bridges, and PN and MN ruptures are largely unexplored in human cancer tissues, as are their underlying causes and *in vivo* consequences.

This uncertainty persists because our current understanding of the process of nuclear rupture derives primarily from live-cell imaging and immunofluorescence (IF) of cell lines *in vitro,* with imaging of only a limited number of human tumor tissues. By contrast, most data on nuclear atypia in human cancers derives primarily from classical histopathologic studies using hematoxylin and eosin (H&E) stained tissues and single-marker immunohistochemistry (IHC), typically with limited associated mechanistic analysis. Studies that bridge these two approaches are largely missing, but this task has been facilitated by the recent development of high-plex, high-resolution IF imaging techniques for the study of human tissues. These high-plex imaging methods, which include the cyclic immunofluorescence (CyCIF) approach used here, make it possible to characterize cell states and nuclear morphology in formalin-fixed paraffin-embedded (FFPE) human tumor specimens that are routinely available in pathology archives.^23^ Furthermore, high-plex imaging enables simultaneous differentiation of cell type (stromal, immune, tumor) and interrogation of functional cell states in a preserved tissue environment. We have previously used such methods to characterize and integrate cellular phenotypes into 2D and 3D tumor atlases as part of the Human Tumor Atlas Network (HTAN).^24–27^

Here, we leverage conventional and high-plex approaches to explore the significance of *sub-cellular* phenotypes such as nuclear atypia and nuclear rupture in glioblastoma (GBM), the most common and lethal adult brain tumor, with a median survival of less than two years.^28,29^ We use a combination of conventional tissue imaging, 3D scanning electron microscopy (EM), and high-plex, high-resolution CyCIF in 2D and 3D, to characterize and quantify NE ruptures and associated phenotypes, focusing on primary human GBM resections and patient-derived xenografts (PDX). We complement these methods with live-cell imaging of the same phenomena in patient-derived cell line models (**Figure 1A**) and analysis of tumor cell states using single-cell RNA sequencing (scRNA-seq) data. This integrated approach allowed us to identify molecular and structural features associated with NE rupture in glioblastoma tissues and demonstrate relationships between nuclear envelope rupture, cell-state differentiation, and innate immune signaling.

**Figure 1:**
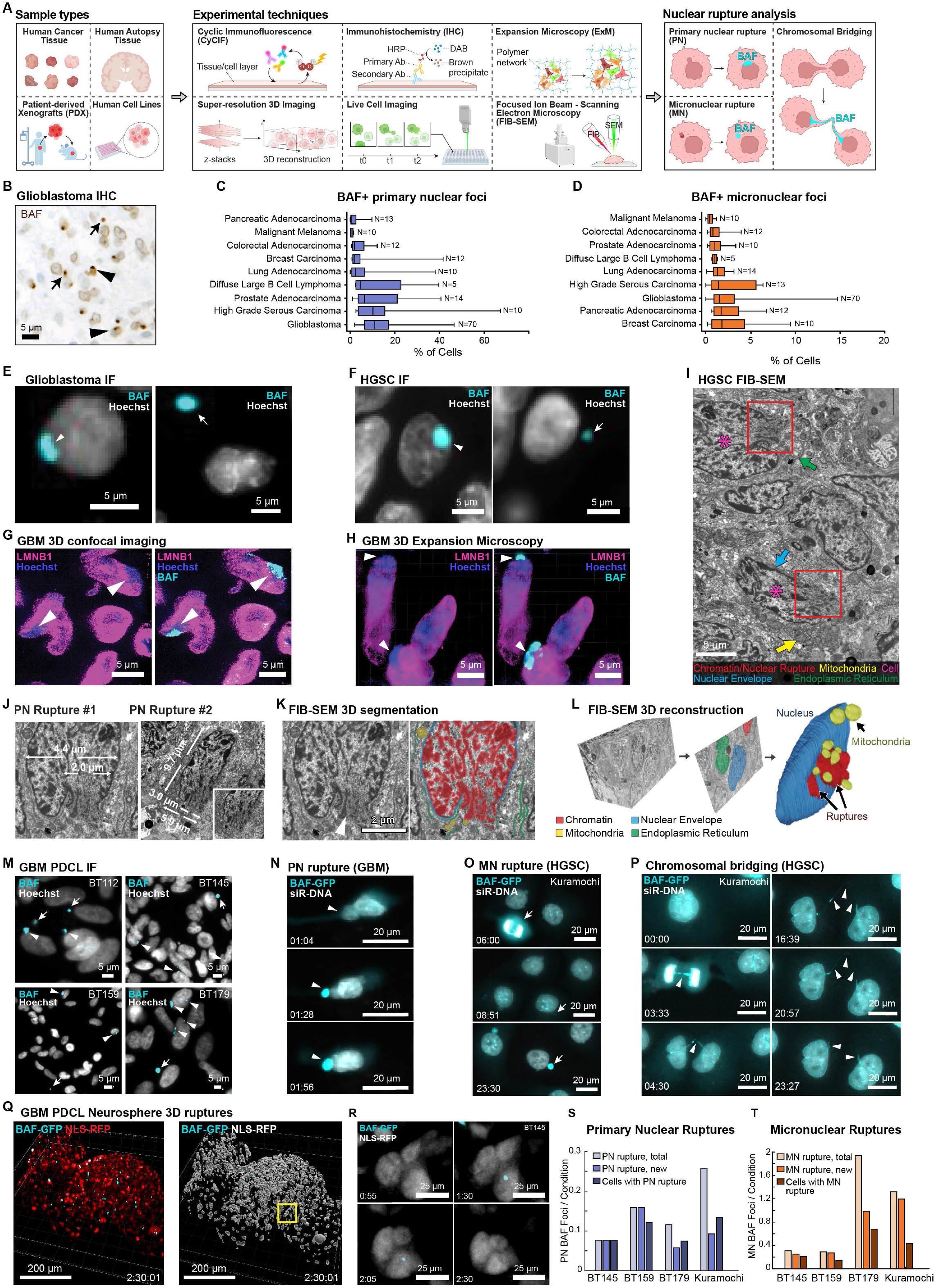
Primary and micronuclear ruptures are common in human cancer. **A,** Schematic of sample types, experimental methods, and features analyzed in this study (generated with BioRender). **B,** Representative BAF immunohistochemistry of human GBM tissue (arrowheads, PN ruptures; arrows, MN ruptures). **C,D,** Quantification of nuclear rupture across multiple cancer types (n=156) showing frequency of PN **(C)** and MN **(D)** rupture (boxes 1st - 3rd quartile, bars 1.5xIQR). **E,F,** Immunofluorescence images of BAF-positive PN ruptures (left panels) and MN ruptures (right panels) in **(E)** GBM and **(F)** high-grade serous ovarian carcinoma (HGSC). **G,H,** 3D confocal (**G**) and 3D expansion immunofluorescence (**H**) showing lamin B1 defects and chromatin herniation indicative of PN rupture at BAF foci in GBM tissue. **I,** Focused ion beam scanning electron microscopy (FIB-SEM) image of human HGSC tissue, showing two atypical tumor cells with NE ruptures (boxes). **J,** Ultrastructural measurement of PN ruptures from two tumor cells. **K,** Representative segmentation of FIB-SEM images. **L,** 3D reconstruction of the FIB-SEM dataset showing nucleus, mitochondria, and herniated chromatin from two PN ruptures interdigitating between mitochondria. **M,** BAF immunofluorescence in unaltered GBM PDCLs showing spontaneous rupture events (arrows, MN rupture; arrowheads, PN rupture). **N-P,** Time lapse microscopy of BAF-eGFP transgenic PDCL labeled with siR-DNA. **(N)** PN rupture associated with nuclear deformation in GBM PDCL (BT145). **O,P,** MN formation and rupture **(O)** and chromosomal bridging (**P**) in Kuramochi ovarian cancer cells following mitotic chromosomal mis-segregation. **Q,** 3D reconstruction and surface rendering of a BAF-eGFP / NLS-RFP transgenic GBM PDCL neurosphere. **R**, Time-lapse imaging of a *de novo* PN rupture event in 3D neurosphere culture, showing formation and resolution of a PN BAF focus. **S**,**T**, Quantification of existing, new, and total ruptures in 2D adherent culture of BAF-eGFP cell lines, for PN (**S**) and MN (**T**). Times shown in hh:mm.

## Results

### Primary and micronuclear ruptures are common in human cancer

The most efficient way to profile panels of human tumors for cell-intrinsic defects such as nuclear rupture is to perform IHC on tissue microarrays (TMAs), each of which can contain multiple samples from dozens of human cancer specimens. Using TMAs, we estimated the frequency of NE rupture in nine cancer subtypes representing most major organ systems and developmental lineages by performing BAF IHC on glioblastoma (**Figure 1B**), lymphoma, melanoma, and carcinomas of the ovary, prostate, lung, breast, colon, and pancreas (total of n=156 primary untreated tumors) (**Figure S1A** and **Table S1**). A board-certified pathologist visually quantified the proportion of tumor cells with BAF foci, which are indicative of NE rupture.^22,30^ BAF foci were found in specimens from all cancer types examined. Ruptures localized to the primary NE were classified as primary nuclear (PN) ruptures (median prevalence 6.3% of cells, range 0-67.3%) (**Figure 1C**), while foci localized to the cytoplasm, typically in a perinuclear distribution, were classified as micronuclear (MN) ruptures (median 1.2% of cells, range 0-14.5%) (**Figure 1D**). Immunofluorescence imaging of human tissue microarrays (TMAs) similarly demonstrated a high frequency of PN and MN BAF foci in GBM (n=145 total tumors) (**Figure 1E**) and also in high-grade serous (ovarian) carcinoma (HGSC) (n=55 tumors) (**Figure 1F**), indicative of NE rupture (**Figures S1B-S1D**).

Next, we confirmed that BAF foci indeed represent nuclear rupture events. First, we characterized the morphology of these BAF foci and nuclear lamina proteins in selected GBM tissues using 3D confocal IF imaging (**Figure 1G** and **Figure S1E**) and 3D expansion-microscopy^31^ (**Figure 1H** and **Figure S1F**). We found that large PN BAF foci were associated with focal lamin B1 defects and chromatin herniation, consistent with NE rupture (**Figures 1G** and **1H)**. To assess the extent of chromatin herniation and cytoplasmic infiltration, we performed serial Focused Ion Beam Scanning Electron Microscopy (FIB-SEM)^32^ on a 35×27×12µm region of a metastatic HGSC (n=43 cells) (**Figures 1I-1L** and **Figure S1G**). Analysis of the electron micrographs revealed multiple highly irregular tumor cell nuclei exhibiting PN rupture with herniation of osmophilic heterochromatin into the cytoplasm (**Figures 1I-1K);** these herniations reached sizes of up to 2-3µm in diameter, as compared to a nuclear diameter of ∼10µm (**Figure 1J**). Detailed segmentation and 3D-reconstruction of nuclei, cytoplasmic organelles (mitochondria, endoplasmic reticulum), and ruptured chromatin showed that chromatin that had herniated into the cytoplasm was often interdigitated between mitochondria and endoplasmic reticulum tubules (**Figures 1K** and **1L**). Moreover, structures appearing in light microscopy as bright BAF foci and large PN ruptures were shown by EM to consist of multiple smaller rupture events in close approximation, indicative of repeated rupture or localized fragmentation of the NE.

Whether cells exhibit similar rates of rupture in cell culture and human tissues is unclear. The temporal dynamics of NE rupture and BAF localization are incompletely characterized, even *in vitro*. We therefore cultured a series of GBM patient-derived cell lines (PDCL)^33^ as 2D monolayers and 3D organoids to quantify rupture frequency and dynamics. Immunofluorescence performed on 2D adherent cultures showed numerous BAF foci including both PN and MN ruptures in unperturbed cells (**Figure 1M**). To study the dynamics of these events by live-cell imaging we introduced a BAF-eGFP reporter into three GBM PDCL cell lines (BT145, BT159, BT179) and the Kuramochi cell line, which has previously been shown to be representative of human HGSC.^34^ DNA was visualized using Silicon Rhodamine DNA dye (SiR-DNA),^35^ and cells were imaged in 2D adherent culture (**Figures 1N-1P** and **Figures S1H-S1J** and **Video S1**). Selected cell lines underwent transduction of NLS-RFP and were imaged using time-lapse confocal microscopy in 3D neurosphere culture showing similar NE rupture events (**Figures 1Q** and **1R** and **Video S1**). Fixed time-point live imaging of 2D adherent cultures showed similar rates of PN rupture compared to human tumor tissues (11.7%; range 7.7-11.6%); an even higher rate was observed in Kuramochi ovarian cells (25.8%) (**Figure 1S**). GBM PDCL and Kuramochi cells also exhibited ongoing MN formation from lagging chromosomes (**Figure 1O** and **Figure S1I**). Interestingly, analysis of MN rupture rate over the full imaging period (24 hours) showed that a majority of GBM cells exhibited at least one MN rupture event (mean 79.6% of cells; range 64.5-94.3%), with similarly high rates in Kuramochi (83.7%), with some cells exhibiting multiple events (**Figure 1T**). These data show that the formation of PN and MN ruptures is ongoing in dividing GBM and ovarian cancer cells.

PN rupture was associated with prominent distortion and blebbing of the primary nucleus in both stationary and motile cells accompanied by compromised primary nuclear envelopes, leakage of DNA into the cytoplasmic compartment (indicated by SiR-DNA), and the formation of intense BAF-eGFP foci. These findings suggest the involvement of both mechanically-induced and spontaneous rupture mechanisms (**Figure 1N** and **Figures S1H** and **S1I**).^36^ BAF-eGFP live-cell imaging also revealed formation and collapse of chromosomal nuclear bridges connecting daughter cells, resulting from incomplete separation of sister chromatids during mitosis^37^ (**Figure 1P** and **Figure S1J**). Time-course analysis of BAF focus intensity in Kuramochi cells and GBM PDCL showed that PN and MN rupture events were transient with both types of rupture resolving over time (**Figures S1K-S1N**). PN ruptures were quicker to resolve, reducing BAF focus intensity (mean range: 59-110 minutes) more quickly than MN ruptures (mean range: 141-476 minutes) (**Table S2**). The duration of heightened BAF signal at rupture sites correlated directly with the area (p=0.003; R^2^=0.5153) (**Figures S1O** and **S1P**) and the intensity (p=0.047; R^2^=0.36) (**Figures S1Q** and **S1R**) of the focus, suggesting that larger ruptures induce greater BAF accumulation and persist longer. Moreover, these data show that quantification of ruptures in fixed cell images, such as those of human cancer tissue, likely underestimates the frequency of the highly dynamic rupture-resolution process.

Collectively, these cross-validated cell line and human tissue specimen data show that susceptibility to NE rupture is a surprisingly common feature of many human cancers. Certain cancer types, such as GBM and HGSC, exhibit particularly high rates of PN and MN rupture, suggesting that these subtypes are especially prone to nuclear instability.

### Multiplexed subcellular imaging of cell line models reveals primary and micronuclear rupture are associated with distinctive protein expression patterns

To develop a robust imaging assay to quantify the relationship between NE rupture and cell state, we developed a multiplex antibody panel for plate-based cyclic-immunofluorescence (p-CyCIF)^38^ of hTERT-immortalized retinal pigmented epithelium (RPE) cells subjected to diverse genetic, chemical, and environmental perturbations designed to alter NE function and structure. RPE cells are a common experimental model for studying nuclear envelope biology *in vitro*.^39^ Perturbations included: (**i**) *BANF1* siRNA (BAF-kd) as a negative control for focus formation, (**ii**) *LMNB1* siRNA (LMNB1-kd) to induce spontaneous PN rupture,^40^ (**iii)** NMS-P715, a MPS1 kinase inhibitor (MPS1i), to induce micronucleus formation,^10^ (**iv**) X-ray radiation (XR) to induce MN formation and nuclear atypia,^41^ or (**v**) exogenous herring testis (HT) DNA transfection to induce STING focus formation (**Figure 2A**). Using p-CyCIF, we examined the subcellular localization of multiple proteins including BAF, nuclear lamina proteins (Lamin A/C, B1, B2), LEM-domain family proteins associated with the nuclear lamina (Lamin B Receptor (LBR) and Emerin), the nuclear pore complex protein NUP133, the DNA damage marker phospho-H2AX (pH2AX), and innate immune signaling proteins (cGAS, STING), which can be activated when DNA is released into the cytoplasm (see **Table S3**).^19,20^

**Figure 2:**
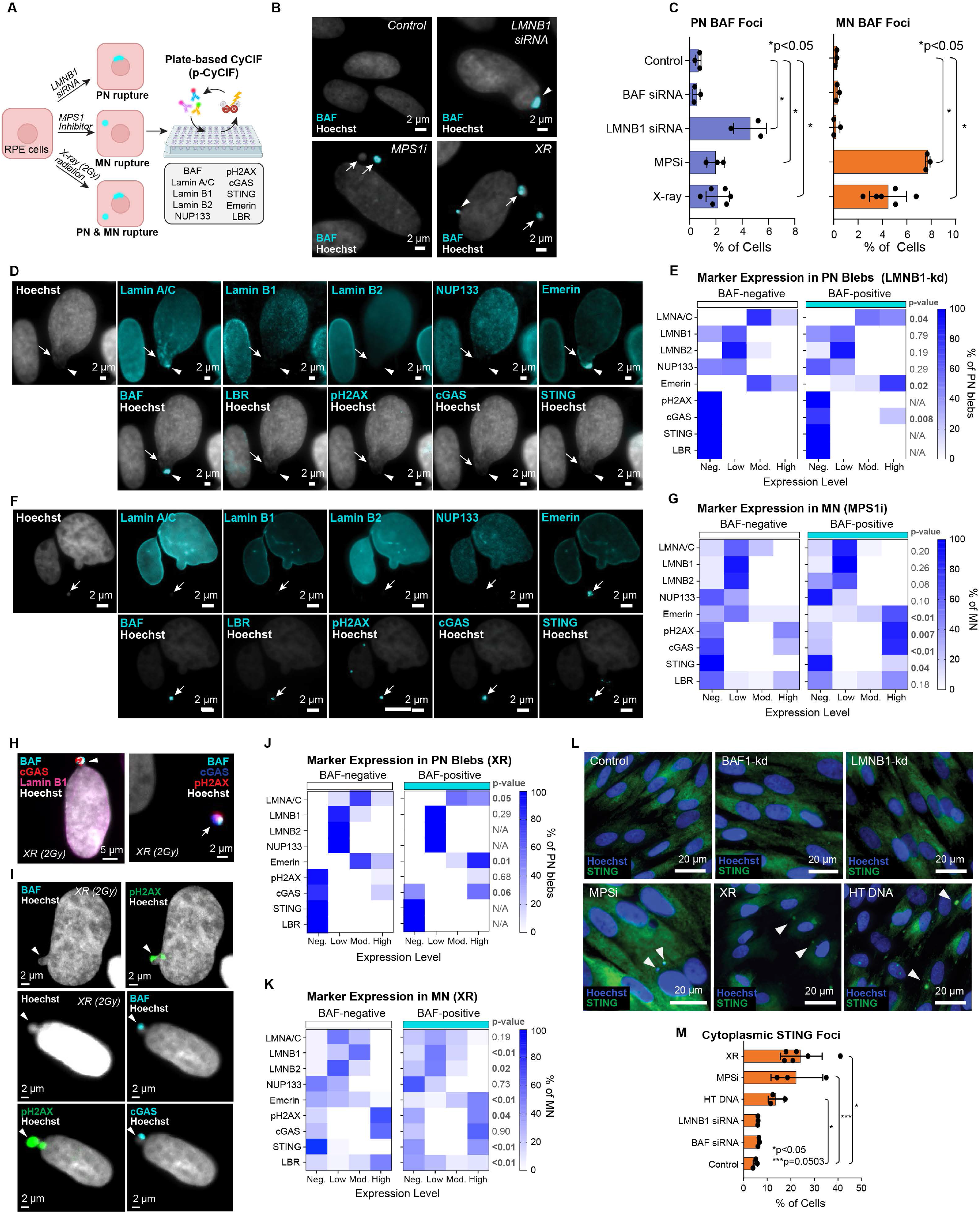
Multiplexed subcellular imaging of cell line models reveals primary and micronuclear rupture are associated with distinctive protein expression patterns. **A,** Experimental overview. Human RPE cells were treated with *LMNB1* siRNA, MPS1i, or X-ray radiation (2Gy), to induce PN or MN rupture then analyzed using plate-based CyCIF. **B,** Representative images of each treatment group (arrows, MN rupture; arrowheads, PN rupture). **C,** Quantification of BAF-positive PN and MN ruptures following treatment (mean % of cells +/- S.D) (*p<0.05) (t-test, two-sided, unpaired). **D,** p-CyCIF images of a ruptured, BAF-positive PN bleb in cells treated with *LMNB1* siRNA, showing localized disorganization and accumulation of lamin A/C and emerin, reduced levels of lamin B1/B2 and NUP133, and absence of pH2AX, LBR, and STING. **E,** Percentage of BAF-negative or BAF-positive PN blebs in *LMNB1*-kd cells expressing each intensity of a given marker, based on semi-quantitative visual scoring (t-test, two-sided, unpaired). **F,** p-CyCIF images of a ruptured, BAF-positive micronucleus a cell treated with MPS1 inhibitor, exhibiting low lamin A/C, lamin B1/B2, and NUP133, and accumulation of emerin, LBR, pH2AX, cGAS, and STING. **G,** Percentage of BAF-negative or BAF-positive MN expressing each intensity of a given marker in MPS1i-treated cells, based on semi-quantitative visual scoring (t-test, two-sided, unpaired). **H,** Representative images of XR-treated cells with PN and MN ruptures associated with accumulation of cGAS and/or pH2AX. **I,** Representative images of XR-treated cells with occasional pH2AX-positive PN blebs. **J,K,** Percentage of BAF-negative or BAF-positive PN (J) or MN (K) with a given expression level of indicated markers. **L,** Representative p-CyCIF images of STING in RPE cells per condition (arrowheads, STING foci). **M,** Quantification STING foci in each experimental condition (*p<0.05, ***p=0.0503) (t-test, two-sided, unpaired).

In control RPE cells, nuclear blebs (protrusions of the nuclear envelope) and PN BAF foci were infrequent (mean 0.63% of cells/well, n=2,365) and were reduced by *BANF1* knockdown (0.53%, n=3034) (**Figures 2B** and **2C**; **Table S4**). *LMNB1*-kd cells exhibited increased blebbing and a 7.0-fold increase in the prevalence of PN BAF foci (4.4%, n=2,672; p=0.005) (t-test, unpaired, two-sided), consistent with higher rates of PN rupture. Cells treated with MPS1i (2%; 3.2-fold; p=0.03) or XR (2.4%; 3.8-fold; p=0.03) also exhibited modestly increased PN rupture. MN BAF foci were rare in control cells (mean 0.2% cells/well) and BAF-kd cells (0.3%). However, treatment with the MPS1i resulted in a 35.6-fold increase in MN BAF foci (7.5%, n=1170 cells; p=3.3×10^-7^), consistent with increased rates of MN rupture. Exposure to XR (n=615) also significantly increased MN BAF foci (5.0%; 23.8-fold; p=0.002). Thus, factors that promote PN and MN rupture are similar but not identical.

Visual analysis of p-CyCIF data from LMNB1-kd, MPS1i, and XR-treated cells (n=214 total events) (**Table S5**) showed that blebs were often associated with focal irregularities in the A-type lamina proteins lamin A/C and emerin, coincident with reduction in the B-type lamina proteins lamin B1 and B2 and nucleoporins (NUP133) (**Figure 2D**). Relative to unruptured blebs, BAF-positive blebs in *LMNB1*-kd cells more often showed higher levels of lamin A/C, emerin, and 23% of these exhibited cGAS accumulation (**Figure 2E**). Both types of blebs typically had low levels of lamin B1 and NUP133 relative to the originating primary nucleus. The DNA damage marker pH2AX, the innate immune effector STING, and LBR were not found to be associated with ruptured or unruptured blebs in *LMNB1*-kd cells. In MPS1i cells, both ruptured and unruptured MN displayed low levels of lamin A/C, lamin B1, lamin B2, and NUP133 consistent with aberrant NE assembly. However, relative to unruptured MN, BAF-positive ruptured MN more frequently showed high levels of emerin, pH2AX (81% of BAF+ MN vs. 46% BAF-), cGAS (78% of BAF+ MN vs. 27% BAF-), and LBR (**Figure 2G**). STING was occasionally observed to localize to ruptured MN (15% of BAF+ MN) and STING+ MN showed cGAS accumulation in every instance (**Figures 2F** and **2G**). This phenomenon was not observed in PN ruptures or unruptured MN. Evaluation of XR-treated cells yielded similar results, with frequent PN and MN ruptures more often showing high levels of cGAS (56% PN, 78% MN), higher levels of lamin A/C and emerin, and lower levels of lamin B1, B2, and NUP133 (**Figures 2H-K**). As with other conditions, STING co-localization with BAF was only observed in a subset of ruptured MN (**Figure 2K**). In XR-treated cells, elevated levels of the DNA damage marker pH2AX were observed in occasional PN blebs that were either ruptured (∼6%) or unruptured (∼11%) (**Figure 2I**). This contrasts with data from *LMNB1-*kd cells in which pH2AX was not associated with PN rupture but may represent an incidental association of XR-damaged DNA with PN blebs in a small subset of cells.

Evaluation of STING staining revealed increased cytoplasmic focus formation, indicative of activation, in cells transfected with HT DNA as compared to control conditions (2.6-fold; 13% of cells; p=0.01) (**Figures 2L** and **2M**). XR (5.8-fold; 28.9%; p=0.01) and MPS1i (4.0-fold; 20.2%; p=0.05) also significantly induced the formation of STING foci, presumably in part due to substantially increased numbers of ruptured micronuclei in these specimens. Neither *LMNB1* siRNA nor *BANF1* siRNA transduction led to significant increases in STING focus formation.

These analyses show that PN and MN exhibit distinctive changes in NE protein levels and evidence of innate immune activation through the cGAS-STING pathway. PN blebs exhibit local reductions in B-type lamina-associated proteins, with disorganization and focal accumulation of A-type lamina proteins such as emerin, and lamin A/C, particularly in ruptured BAF-positive blebs. BAF may recruit A-type lamina proteins to blebs via its LEM binding domains. MN exhibit low levels of lamin A/C, B1, B2, and NUP133; however, ruptured MN often exhibit high levels of emerin and LBR, the latter of which is not seen in PN rupture. Notably, both ruptured and unruptured MN were associated with accumulation of the DNA damage marker pH2AX, with significantly greater damage in ruptured MN. pH2AX staining was not observed in PN blebs except in a small subset of irradiated cells in which damage may be caused by XR-treatment rather than rupture itself. XR treatment and MPS1 inhibition each induced MN formation and rupture, leading to significantly increased STING focus formation, but this effect was not seen in cells with PN rupture, such as *LMNB1-*kd cells. These data provide the first multiplexed, high-dimensional profiling of individual blebs and MN, including both ruptured and unruptured structures, directly illustrating that multiple markers are simultaneously altered in atypical nuclear structures undergoing NE rupture, with related but distinct phenotypes for PN and MN rupture.

### Nuclear envelope rupture is associated with differences in lamin subtype expression in glioblastoma tissues

One factor that may contribute to nuclear instability in cancer cells is reduced expression of structural components of the nuclear envelope; these include *LMNA* and its spliceoforms lamin A/C^37^, *LMNB1,* and *LMNB2*.^42^ Frequent nuclear rupture and fragmentation is observed, for example, in patients with laminopathy syndromes caused by mutations in *LMNA, EMD* (emerin), and other proteins that make up the nuclear envelope.^43^ To study this association further, we analyzed data from The Cancer Genome Atlas (TCGA) Pan-Cancer Study (n=10,071 tumors from 30 cancer types)^44^, and found that GBM has one of the lowest levels of *LMNA* expression of all cancer types (49.5% of mean of all subtypes; p=1.25×10^-56^), but expresses *LMNB1* at a relatively high level (121% of mean, p=2.90×10^-4^) and shows moderate expression of *LMNB2* (86.5% of mean, p=1.20×10^-3^) (**Figures S2A-S2C**). Low levels of *LMNA* expression were not associated with measurable genomic aberrations in the *LMNA* gene (**Figures S2D** and **S2E**). Analysis of RNAi and CRISPR knockout data from The Cancer Dependency Map (DepMap)^45^ showed that GBM and diffuse glioma cell lines are the most dependent on *LMNA* expression as compared to all other cancers in the CCLE (n=1,095 lines; p=1.43×10^-20^ and 9.2×10^-25^ respectively for CRISPR knockouts) with lines derived from mesothelioma and hematopoietic malignancies also showing significant dependence (**Figures S2F** and **S2G**). Glioma lines were also dependent on *LMNB1*, albeit with a weaker effect (p=4.06×10^-6^) (**Figures S2H** and **S2I**), whereas *LMNB2* showed no strong dependence in any cell line (**Figures S2J** and **S2K**). Thus, GBM and glioma cell lines express low levels of *LMNA* and are dependent on *LMNA* and *LMNB1* for survival.

To study lamin expression in human GBM we performed 28-plex t-CyCIF on a glioma TMA comprising 145 GBM tissues, including 127 IDH-WT GBM (82 primary, 45 recurrent; this type of GBM represents ∼90% of cases) and 15 IDH-mutant GBM (4 primary, 11 recurrent) (**Figure 3A**). The CyCIF panel included antibodies against lamins validated using p-CyCIF (**Figure 2**), along with antibodies against proteins involved in NE rupture and repair, DNA damage, proliferation, and cell identity. Inspection of TMA cores from 82 untreated primary IDH-WT GBM revealed substantial variation in lamin protein levels across tumors and among cells in the same tumor, with an apparent inverse correlation between lamin A/C levels and PN rupture (**Figure 3B**). Processing the images to segment individual cells from the cores yielded ∼1.6 x 10^5^ SOX2-positive tumor cells (total cells= 2.6 x 10^5^) among which 1.3 x10^4^ (7.8%) had micronuclei, 644 had BAF+ MN (0.5%) and 5.9 x10^4^ had BAF+ PN (34%) (**Figures 3C** and **3D**). Quantitative single-nucleus analysis of A-type (A/C) and B-type (B1 and B2) lamin levels in all SOX2-positive tumor cells confirmed a strong inverse correlation between PN lamin A/C and the presence of a PN rupture (p=4.1×10^-37^), suggesting that low lamin A/C levels render tumor cells susceptible to PN rupture. In contrast, there was no strong correlation between PN rupture and lamin B2, while lamin B1 levels were higher in cells with PN ruptures (p<1×10^-40^) (t-test, two-sided, unpaired) (**Figure 3E**). We found that ruptured MN were associated with significantly lower levels of lamin B1 and B2, with no strong difference in lamin A/C compared to unruptured MN (**Figure 3F**). Thus, at a single-cell level, PN and MN rupture frequencies were strongly correlated with differences in underlying lamin expression levels: low A-type lamins for PN and low B-type lamins for MN.

**Figure 3:**
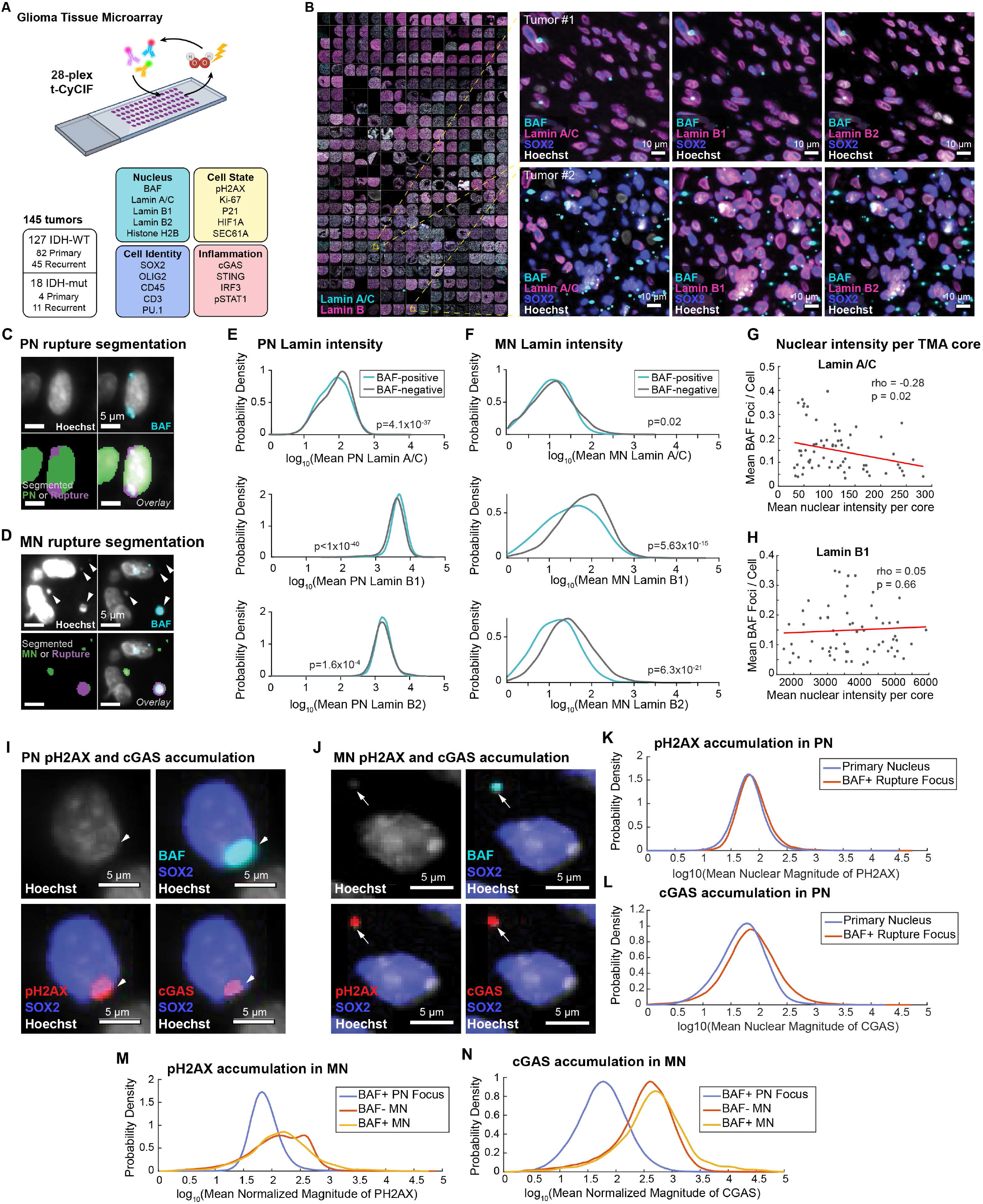
Nuclear envelope rupture is associated with differences in lamin subtype expression in glioblastoma tissues. **A,** Overview of 28-plex t-CyCIF experiment on a glioma tissue microarray (TMA). **B,** Selected t-CyCIF images of glioma TMA with cores de-arrayed. Right top: primary IDH-WT GBM tumor with high lamin A/C expression and few PN ruptures. Right bottom: primary IDH-WT GBM tumor with low lamin A/C expression and many PN ruptures. **C,D,** Examples of segmentation of PN rupture (**C**) and MN rupture (**D**). **E,F,** Probability density plots of lamin A/C, lamin B1, and lamin B2 protein expression in 161,834 PN (**E**) and 5,974 MN (**F**) from 82 primary IDH-WT GBM. P values as indicated (t-test, two-sided, unpaired). **G,H,** Scatter plot of the mean number of BAF foci per cell versus mean nuclear lamin A/C or lamin B1 expression in primary IDH-WT GBM (N=82, duplicate cores). Pearson correlation analysis shows that lamin A/C is inversely correlated with PN BAF focus frequency (rho=-0.28, p=0.02) (**G**); lamin B1 shows no significant association (rho=0.05, p=0.66) (**H**). **I,J,** t-CyCIF images of BAF, pH2AX, cGAS in primary IDH-WT GBM with PN (**I**) and MN (**J**) ruptures. **K,L,** Probability density plots of pH2AX (**K**) and cGAS (**L**) signal in segmented BAF+ rupture foci and BAF-negative PN. **M,N,** Probability density plots of pH2AX (**M**) and cGAS (**N**) signal in segmented BAF-positive PN and BAF-negative and -positive MN.

To assess this effect at a population level we quantified the number of PN BAF foci compared to the mean nuclear intensity of each lamin protein across all tumor cells in each core. We again found that PN lamin A/C staining was inversely correlated with PN rupture frequency across all tumors in SOX2 positive tumor cells (ρ=-0.28, p=0.021) (**Figure 3G**); no relationship with lamin B1 levels and PN rupture was observed (ρ=0.05, p=0.66) (**Figure 3H**). When we compared primary IDH-WT GBM (n=82) to recurrent/residual tumors post-chemoradiation (n=45) we found that tumor cells in recurrent cases expressed similar levels of lamin B1 and B2, slightly higher lamin A/C, and reduced rupture rates, though this effect did not reach statistical significance, possibly due to the smaller set of recurrent tumors (p=0.157) (**Figure S2L**). Comparison of IDH-WT GBM to IDH-mutant GBM showed similar levels of lamin B1 and B2, but reduced lamin A/C (p<0.0001) in IDH-mutant cases, with a corresponding increase in PN rupture rate (p<0.0001) (**Figure S2M**). Thus, CyCIF data show that variation in lamin protein expression typically correlates with NE rupture frequency.

CyCIF data additionally showed that PN (**Figure 3I**) and MN (**Figure 3J**) ruptures in primary untreated GBM tissues can show focal expression of the DNA damage marker pH2AX and innate immune marker cGAS. This contrasts with data from RPE cells, in which DNA damage was not associated with PN ruptures except in XR-treated cells. Quantitative analysis of tissue CyCIF data at the level of individual PN and MN BAF foci revealed a continuous Gaussian distribution of pH2AX and cGAS signals across all data points, suggesting that rupture events are associated with varying levels of DNA damage and innate immune activation. We found that mean pH2AX levels were only modestly increased in PN ruptures relative to the originating primary nuclei (∼1.3-fold; p=1.8×10^-4^) (**Figure 3K**), whereas mean cGAS levels were substantially higher at PN rupture sites compared to the originating primary nucleus (∼3.3-fold; p=4.3×10^-54^) (**Figure 3L**). Analysis of pH2AX levels associated with BAF-positive MN (normalized to DNA content) showed that mean expression was significantly higher (2.3-fold, p<1×10^-40^) than in PN rupture events (**Figure 3M**). Similarly, cGAS was markedly enriched in MN ruptures relative to PN ruptures (5.7-fold, p<1×10^-40^) (**Figure 3N**). Ruptured MN showed 50% greater mean pH2AX expression than unruptured MN (p=1.46×10^-20^). cGAS was also strongly elevated in ruptured MN compared to unruptured MN (2.4-fold; p<1×10^-40^). Thus, while PN ruptures show significant accumulation of cGAS, PN rupture is associated with limited DNA damage in untreated human cancer tissues, while MN are associated with dramatically increased levels of DNA damage and cGAS accumulation, particularly in those with NE rupture.

### Primary nuclear rupture and lamin expression are correlated with GBM tumor cell states

Single-cell RNA sequencing (scRNA-seq) studies have shown that GBM cells predominantly exist in four molecular states defined by transcriptional signatures: oligodendrocyte progenitor cell-like (OPC), neural progenitor cell-like (NPC), astrocyte-like (AC), and mesenchymal-like (MES).^46^ To determine whether these states are associated with differences in NE rupture frequencies and lamin protein expression, we analyzed publicly available scRNA-seq data from 28 adult and pediatric GBM specimens, comprising a total of 24,131 cells.^46^ We observed that expression of *LMNA* mRNA was significantly lower in OPC and NPC cell states as compared to AC and MES (mean OPC/NPC *LMNA*=∼57% of mean AC/MES *LMNA*; p<1×10^-16^). Conversely, *LMNB1* expression was significantly higher in OPC and NPC cell states (p<0.0001) (**Figure 4A**).

**Figure 4:**
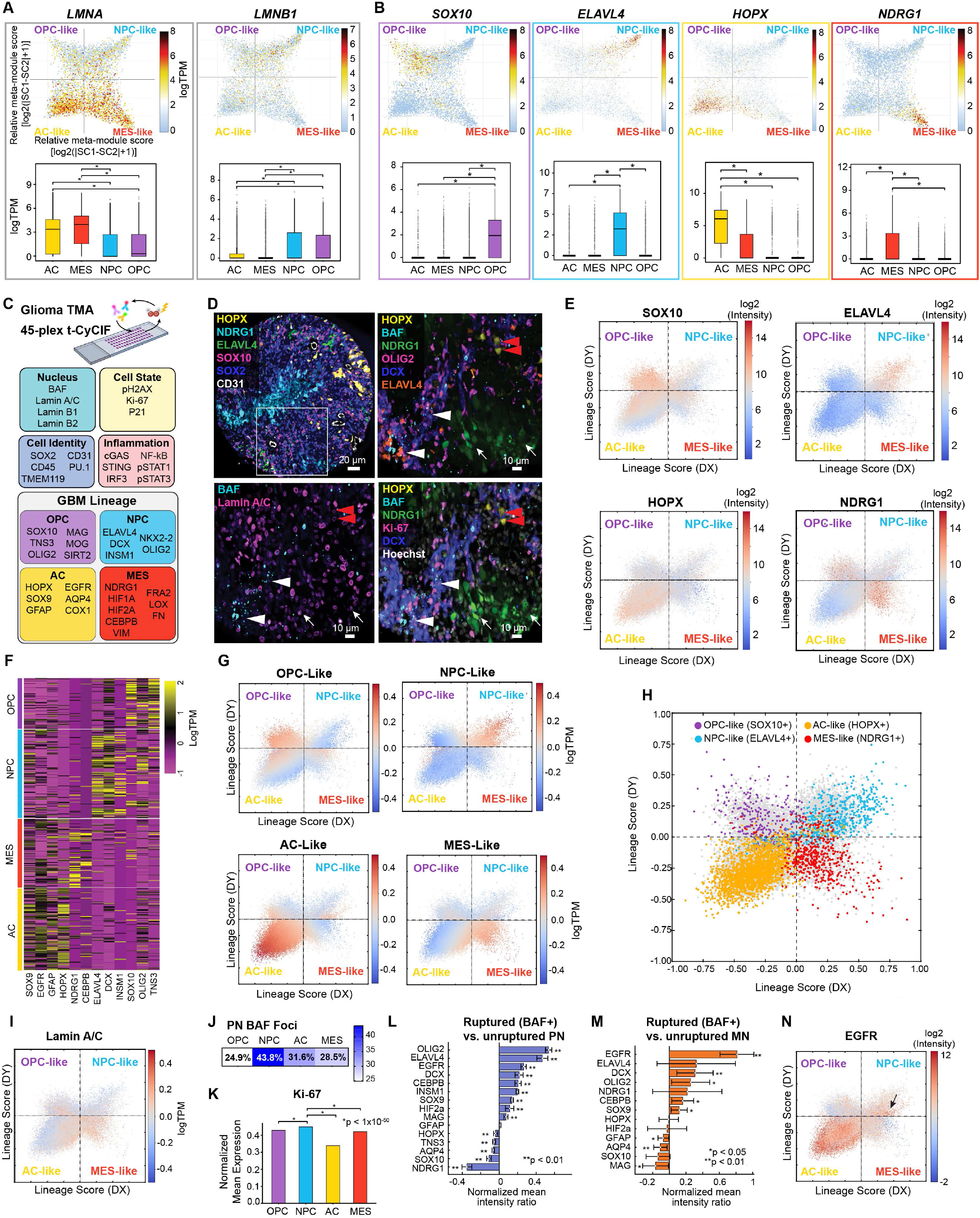
Primary nuclear rupture and lamin expression are correlated with GBM tumor cell states. **A,** scRNA-seq from 28 adult and pediatric GBM (n=24,131 cells) showing lamin RNA expression by GBM cell state. *p<0.05; t-test, two-sided, unpaired. **B,** Plots of scRNA-seq data showing putative markers of each cell state in GBM (*p<1×10^-16^) (t-test, two-sided, unpaired). **C,** Schematic of t-CyCIF experiment and markers to explore rupture and lineage correlations (generated with BioRender). **D,** Representative GBM t-CyCIF images showing cell state marker expression in distinct tumor cell populations, with reduced lamin A/C expression and increased PN rupture in NPC-like cells (DCX, ELAVL4) (white arrowheads) compared with NDRG1-expressing MES-like cells (white arrows) and HOPX-expressing AC-like cells (red arrowheads). TMA core image with ROI indicated (upper left). **E,** Two-dimensional representation of GBM cell states from t-CyCIF. Each quadrant represents a cell state (based on Neftel *et al*.), with position reflecting the relative score and indicated cell state scores for each cell and intensity values for the indicated markers. **F,** Composite representation of cell state groups in scRNA-seq data for validated markers. **G,** Plot of scores for each cell state mapped to individual tumor cells, showing an excellent correspondence of cell state designations at a single-cell level in t-CyCIF data. **H**, Correlation of manual gating of marker intensity and designation of cells according to core cell state markers and mapping of continuous cell state data. **I**, mapping of lamin A/C levels by cell state score shows that NPC-like cells exhibit the lowest lamin A/C expression. **J**, Quantification of PN rupture rate in t-CyCIF TMA data normalized to cell count. **K,** Quantification of mean nuclear Ki-67 expression in different cell states shows significantly higher proliferation rates in NPC-like cells compared to other populations (p<1×10^-50^ for all comparisons). **L,M,** Quantification of cell state markers in 82 primary IDH-WT GBM comparing cells with ruptured and unruptured PN (**L**) and MN (**M**), showing enrichment of NPC cell state markers and EGFR in ruptured cells compared to the mean of all markers (**p<0.01) (t-test, two-sided, unpaired). **N,** two-dimensional mapping of EGFR expression by cell state shows that while AC-like cells exhibit the strongest overall expression, a secondary cluster of NPC-like cells also exhibits EGFR expression.

There are no known transcriptional signatures for PN or MN rupture. We therefore sought to leverage high-plex tissue imaging to study whether NE rupture correlates with cell state differentiation. To accomplish this, we analyzed scRNA-seq data for genes exhibiting lineage-specific expression and selected those genes encoding proteins with available antibodies. This yielded 3 to 8 putative protein markers for each of the four GBM sub-states (**Figure 4B** and **Figures S3A-S3D).** We then developed a 45-plex t-CyCIF antibody panel that leveraged the capabilities of multiplexed imaging to include multiple cell state markers for each lineage, as well as antibodies against nuclear envelope proteins, cell type markers, regulators of innate immune signaling, and cell proliferation markers (**Figure 4C** and **Table S3**). SOX2, a commonly used marker of glioma cells which is broadly expressed by all tumor lineages (**Figure S3E**), was used to identify tumor cells in general.

Visual review of t-CyCIF data from IDH-WT GBM TMA cores (n=82 tumors) focused on comparison of cells exhibiting an OPC-like (SOX10, TNS3, and MAG protein markers), NPC-like (ELAVL4, DCX, and INSM1), AC-like (HOPX, SOX9, GFAP, EGFR, and AQP4) or MES-like state (NDRG1, HIF1A, HIF2A, and CEBPB). Across GBM and glioma TMAs, we found that that most cores contained cells with markers from each of the four putative GBM cell states, with the proportions of the different states varying from core to core, and individual states often clustering together in space (**Figure 4D**). We found that AC-like markers were typically the most abundant, consistent with the pathologic definition of glioblastoma as an astrocytoma. Cells with NPC-like markers were often found in tight clusters, while cells with OPC-like expression were intermixed throughout the tumor. Markers of the MES cell state (which is often associated with tumor hypoxia) were typically found in peri-necrotic tumor cells. To better understand these patterns, for each CyCIF marker, we computed differential protein expression values from log2-transformed image intensities and identified AC, MES, OPC, and NPC-like states from a 2D projection of the hyperdimensional image data (see **Methods** for details). Plotting the expression of individual CyCIF markers as single-cell profiles showed state-specific patterns of protein expression (**Figure 4E**) reminiscent of the patterns observed in the scRNA-seq data (**Figures 4B** and **4F** and **Figures S3A-S3D**). Composite scores comprising the expression of multiple lineage markers also resolved the individual cells into the four cell state phenotypes (**Figure 4G**). To further confirm these results, we reviewed tissue microarray specimens and performed manual gating of cell states using the most lineage-restricted markers (ELAVL4, HOPX, SOX10, NDRG1) with expression thresholds determined visually by a pathologist. Classification of individual cells by manual gating showed excellent correlation with unbiased mapping of states using continuous intensity data (**Figure 4H**).

At the protein level, NPC-like tumor cells exhibited the lowest levels of lamin A/C expression (**Figure 4I**) and had the highest rates of PN rupture (43.8%) when normalized to cell state frequency (**Figure 4J**). The median rupture rates were higher in this analysis compared to visual quantification of IHC-stained human tissues (**Figure 1**), likely because the broad panel of lineage and cell state markers available in CyCIF improved the exclusion of non-tumor populations (which can make up over 50% of a tumor sample). NPC-like cells exhibited significantly higher mean Ki-67/MIB-1 signal (**Figure 4K**) and accordingly higher proliferation rates (∼37% Ki-67 positive; with normalized mean marker intensity > 0.5 scored as “positive”) as compared to 19-30% for the other cell types (p<1×10^-50^ for each comparison). Analysis of single markers also revealed a strong correlation of PN rupture with NPC markers (OLIG2, ELAVL4, DCX and INSM1) across all SOX2+ tumor cells (**Figure 4L**). MN rupture was also more strongly correlated with NPC-like differentiation (**Figure 4M).** EGFR protein expression also correlated strongly with PN and MN rupture, and qualitative inspection of tissues showed that clusters of NPC-like cells with elevated EGFR expression are likely responsible for this trend (**Figure 4N**); a similar cluster of EGFR^high^ NPC-like cells was also present in scRNA-seq data (**Figure S3A**). While *EGFR* amplification and expression are more commonly associated with astrocytic (AC-like) differentiation, these data suggest there is a distinct PN rupture phenotype in EGFR^high^ NPC-like cells. This suggests that AC, MES, OPC, and NPC types can be further stratified into sub-types based on specific signaling or genetic programs, and that composite analysis of multiple markers is likely necessary to precisely define cells according to lineage cell states.

### Multiplexed 3D confocal imaging of nuclear instability and cell state in glioblastoma

To better visualize the relationship between nuclear instability and cell states we performed 30-plex 3D confocal CyCIF on 20-micron thick tissue sections (with 0.076 x 0.076 x 0.25cm voxel dimensions), including markers of nuclear stability, cell state, DNA damage, and immune signaling (**Figure 5A**). Unlike conventional 5-micron thick FFPE sections that typically capture only a partial slice of a cell, 20-micron sections can capture entire cells – a useful feature for characterizing complex subcellular phenomenon such as NE rupture in 3D.

**Figure 5:**
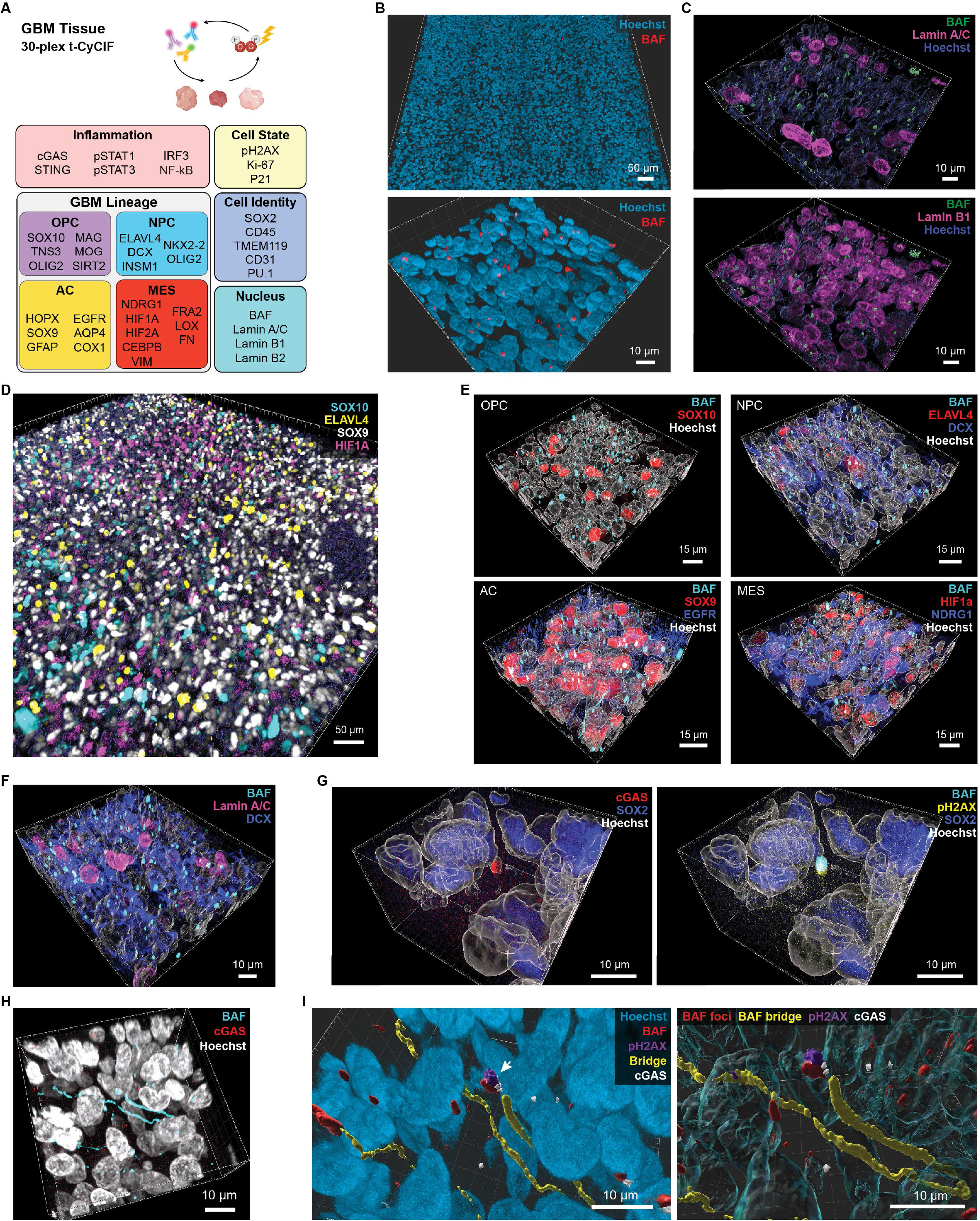
Multiplexed 3D confocal imaging of nuclear instability and cell state in glioblastoma. **A,** Schematic of experimental plan for 3D confocal CyCIF, with indicated markers (generated in BioRender). **B,** High-resolution confocal imaging of GBM tissues reveals nuclear morphology and abundant PN and MN rupture events. **C**, 3D confocal CyCIF confirms that PN ruptures are correlated with reduced expression of lamin A/C, but not lamin B1. **D**, 3D confocal CyCIF of primary IDH-WT GBM showing highly heterogeneous expression of cell state markers **E,** Examination of cell state markers and BAF confirms enrichment of PN BAF foci in NPC-like cells (DCX+, ELAVL4+), compared to other cell state populations. **F,** NPC-like cells exhibit low expression of lamin A/C and high rates of BAF-positive PN rupture. **G,** 3D confocal CyCIF shows colocalization of BAF, pH2AX, and cGAS in a ruptured MN **H,** 3D confocal CyCIF enables visualization and characterization of complex nuclear structures including chromosomal bridges. **I,** 3D reconstruction and mapping of chromosomal bridges reveals their complex architecture, including hairpin turns, and focal ruptures with BAF, pH2AX, and cGAS accumulation.

3D reconstruction of confocal imaging data provided high-resolution views of nuclear morphology and BAF-positive rupture events in large regions of untreated primary human glioblastoma tissue (**Figure 5B** and **Video S2**). This allowed us to confirm that PN rupture is inversely correlated with lamin A/C (but not lamin B1) levels in tumor cells (**Figure 5C**). Mapping of GBM cell states showed intermixed and highly heterogeneous populations (**Figure 5D**). Simultaneous visualization of BAF confirmed that NPC-like cells (DCX+, ELAVL4+) had more PN ruptures compared to other cell states (**Figure 5E**) and had low levels of lamin A/C (**Figure 5F**). 3D confocal imaging also provided specific and detailed confirmation that cGAS and pH2AX localize to ruptured PN and MN in human tumors (**Figure 5G**). An unexpected benefit of high-resolution imaging of intact whole cells was the ability to clearly visualize chromosomal bridges and their winding, non-linear course between cells (including hairpin turns) (**Figure 5H**). These structures result from improper separation of sister chromatids during anaphase and were strikingly common; we detected 52 bridges among the ∼3,370 SOX2+ tumor cells implying that ∼3.1% of tumor cells have bridges (assuming each bridge involves two daughter cells). Many chromosomal bridges extended for over 10µm and had BAF accumulation along their lengths and focal cGAS and pH2AX (**Figure 5I**).

From these data, we conclude that while PN and MN rupture may occur in any GBM cell state, they are found at disproportionate rates in NPC-like cells, which exhibit a distinctive phenotype characterized by low lamin A/C expression. Cell line data suggest that PN rupture is tied to low lamin A/C expression in this population, but higher proliferation rates may underlie the greater correlation of NPC-like cells with MN rupture, since MN formation typically occurs following chromosomal missegration events during mitosis. We also find that chromosomal bridges in tumors can accumulate BAF, cGAS, and pH2AX at multiple locations, reminiscent of the bridge collapse and NE rupture that we observed in live-cell imaging (**Figure 1** and **Video S1**). Bridging is difficult or impossible to characterize by conventional light or IF microscopy of standard 5µm thick tissue sections. However, it is known to drive oncogenic progression via chromothripsis or DNA damage, arguing for further analysis of human tumors using high-resolution 3D imaging of thick specimens.

### Multiplex imaging of whole-slide tissue specimens reveals the spatial organization of GBM tumor cell states

Although TMA cores are an efficient way to study cell-intrinsic features of tumor cells, they do not provide an accurate assessment of spatial relationships within tissues; whole-slide imaging is required.^25^ We therefore performed t-CyCIF on ∼6 cm^2^ whole-slide human autopsy specimens (n=4; ∼2 x 10^6^ cells per specimen vs. 10^3^ cells in a 0.6 mm TMA core) (**Figure 6A**). Given that protein preservation is an issue in autopsy specimens as compared to surgical specimens,^47^ we focused our analysis on antibodies with the highest signal-to-noise ratio across multiple specimens. Visual inspection by a pathologist (**Figure 6B**) revealed distinct regional patterning of cell state markers in SOX2-positive tumor cells. The MES marker NDRG1 and HIF1A were predominantly expressed by peri-necrotic tumor cells (**Figure 6B** and **Figure S4A**). Consistent with our results from TMA imaging experiments (**Figure 3**), GBM cells with NPC differentiation in the autopsy specimens (i.e., ELAVL4 and/or DCX-expressing cells) had lower levels of lamin A/C and higher rates of PN rupture relative to other cell states (**Figure 6C**). This observation was confirmed by quantitative analysis using GBM cell state scores (**see Figure 4, Methods**) in which SOX2+ tumor cells were clustered according to the four major GBM cell states (**Figure 6D**). Spatial mapping of these cell states in whole-slide images (WSI) recapitulated findings from qualitative inspection, typically showing enrichment of MES-like cells in peri-necrotic regions, NPC and/or OPC markers enriched at the invasive edges, and AC-like markers in the tumor core (**Figure 6E** and **Figure S4B**).

**Figure 6:**
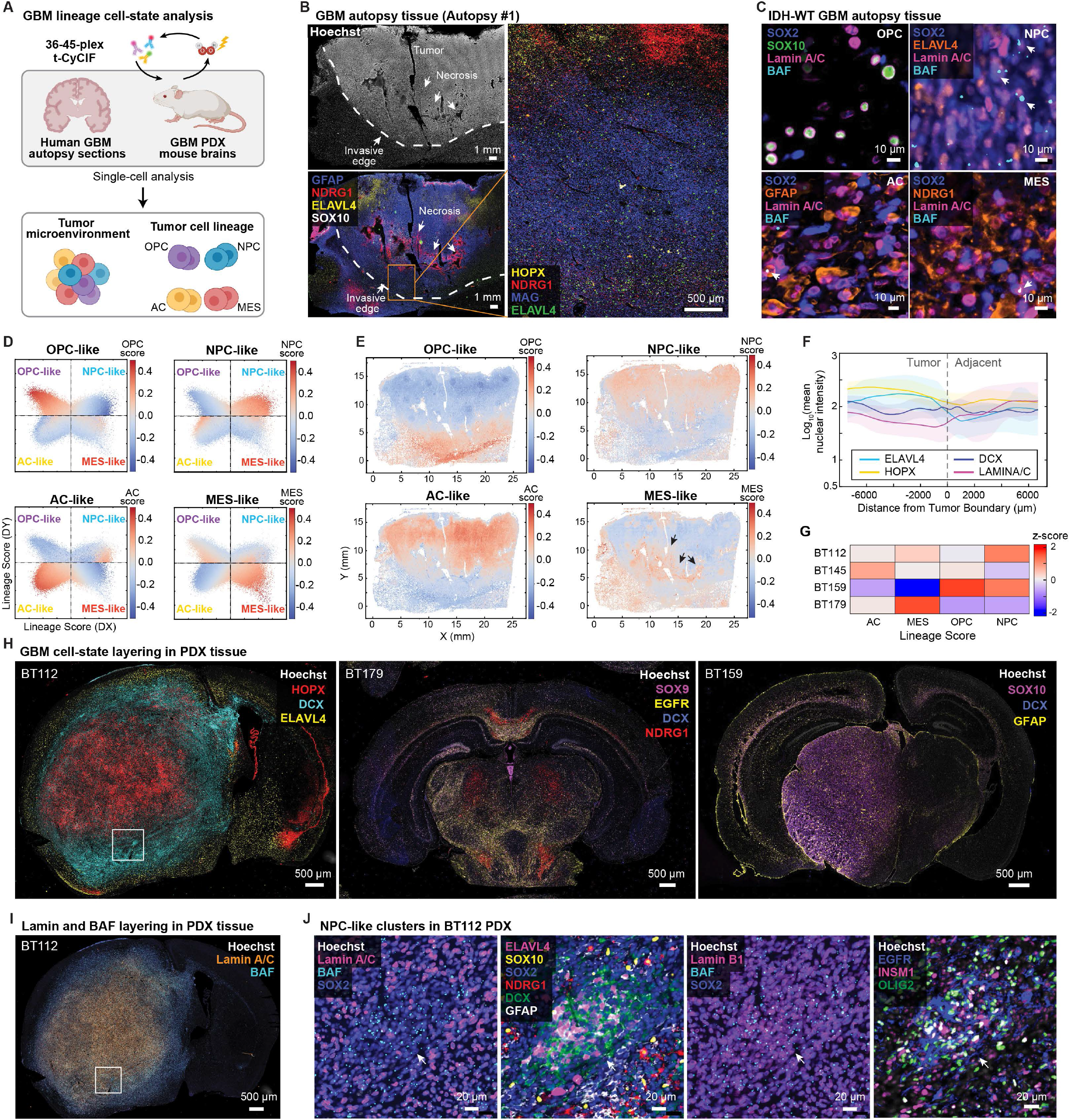
Multiplex imaging of whole-slide tissue specimens reveals the spatial organization of GBM tumor cell states. **A,** Schematic of sample types and design of GBM cell state analysis. **B,** Human GBM autopsy tissue with tumor, necrotic areas, and tumor-brain invasive margin indicated. t-CyCIF images showing representative cell state markers. **C,** Representative t-CyCIF images of human IDH-WT GBM autopsy cell state regions. NPC-like regions exhibit low lamin A/C and higher rates of PN rupture than other cell state regions (arrows, BAF-positive PN ruptures). **D,** Two-dimensional representation of all cells from a human autopsy specimen (#1), correlating state-assignment with underlying cell state scores. **E,** Spatial map of lineage cell state scores across a human autopsy specimen, showing regional differentiation of tumor cells. **F,** Quantitative analysis of cell state scores from the center of the tumor to edge and adjacent regions, showing gradual reduction of AC-like markers (HOPX) and lamin A/C and a gradual increase of NPC-like markers**. G,** Analysis of cell state differentiation patterns in RNA-sequencing data from cultured GBM PDCL lines. **H,** Representative t-CyCIF imaging of patient-derived xenografts (PDX) from 3 PDCL lines: BT112, BT159, and BT179. **I**, t-CyCIF imaging of BT112 shows reduced lamin A/C expression and increased NE rupture at the periphery of the tumor, corresponding to regions of NPC-like differentiation. **J,** Representative t-CyCIF images from the invasive margin of BT112 showing a tight cluster of NPC-like tumor cells (DCX+, ELAVL4+, with low lamin A/C and elevated BAF-positive PN ruptures).

When we measured the spatial distribution of cell state markers from the tumor center to the periphery, we found that states were intermixed with distinctive associations in localized regions, consistent with recurrent spatial patterning. In addition, spatial patterns involving lineage markers were also observed on longer length-scales (>1mm). For example, some tumors showed gradual reduction in AC-like markers (HOPX), accumulation of NPC-like markers (ELAVL4, DCX), and a concomitant reduction of lamin A/C expression from the tumor center to periphery. Others exhibited gradual accumulation of AC-like markers towards the periphery (**Figure 6F** and **Figure S4C**). When we analyzed publicly available spatial transcriptomic data from the IVY Glioblastoma Atlas, which was generated from seven spatially distinct regions from 10 adult GBM specimens^48^, we also found that MES markers (NDRG1, CEBPB) were elevated in regions near necrosis, NPC (ELAVL4) and OPC (SOX10) markers were higher toward the infiltrating edge of the tumor, and lamin A/C was lower towards the infiltrating edge (**Figure S4D**).

To determine whether the spatial patterning of cell lineage and NE rupture is encoded by tumor cell states or driven by microenvironmental features we examined multiple genetically-defined (PDX) tissues. t-CyCIF was performed on whole-brain sections from four patient derived cell lines (PDCLs) grown orthotopically in xenografted NSG-mouse models: BT112, BT145, BT159, BT179. These lines have diverse genetic drivers and exhibit varying patterns of lineage differentiation by RNA analysis, with BT112 exhibiting mixed AC, MES, and NPC-like differentiation, BT145 exhibiting AC, OPC, and NPC-like intermixing, BT159 having OPC and NPC-predominant differentiation, and BT179 being MES-predominant (**Figure 6G**). Because PDX tumors were grown in NSG mice, the TME is immunodeficient; however, PDX models have been widely used to study intracellular phenomena and cell state patterning. As with human autopsies, each patient-derived xenograft (PDX) displayed a distinct pattern of spatially organized growth that appeared to be related to the dominant cell state differentiation state exhibited by the cells prior to xenografting (**Figure 6H** and **Figures S5** and **S6**). For example, the BT112 PDCL showed mixed AC-like, MES-like, and NPC-like differentiation *in vitro* by RNA-Seq analysis, and we found that it adopted a multilayered architecture when grown as a PDX, with AC-like markers expressed in regions near the tumor core and NPC markers expressed by clusters of lamin A/C-low cells at the infiltrative edges (**Figures 6H** and **6I** and **Figure S5A**). BAF-positive PN ruptures were very common in BT112; with particular enrichment in lamin A/C low NPC-like cells located at the infiltrative interface of the tumor (**Figure 6J**).

BT145 PDCL, which showed AC-predominant differentiation *in vitro*, with a minor component of other populations (**Figure S5B**), exhibited an AC-like phenotype with numerous intermixed small clusters of lamin A/C-low NPC-like cells which contained high levels of PN rupture (**Figure S5B**). The BT159 PDCL, which showed a strong predominance of OPC and NPC-like differentiation *in vitro*, retained extensive mixed OPC/NPC differentiation in xenografts with limited expression of AC and MES-like markers (**Figure 6H** and **Figure S6A**). The BT179 PDCL, which had an AC and MES-predominant signature *in vitro*, exhibited diffuse infiltration of the brain stem with AC and MES-like marker expression and limited NPC and OPC-like marker expression in PDX tissue (**Figure 6H** and **Figure S6B**). These data suggest that tumor architecture is self-organizing according to intrinsic differentiation states present in PDCL prior to transplantation that are maintained during tumor initiation and outgrowth. Moreover, given the immunodeficient state of the PDX models, self-organization is not dependent on tumor-immune interactions

Collectively these data confirm that PN rupture correlates with NPC-like tumor cells exhibiting low lamin A/C expression relative to other GBM cell states, and that rupture rates in human autopsy and PDX models are similar to those observed in human resection tissue. At a local level, cell states are often associated with specific anatomic regions; namely, MES-like with peri-necrotic regions, NPC and OPC-like with the infiltrative edges of tumors, and AC-like with the tumor core. However, while tumors with astrocytic-predominant differentiation (e.g., BT112) exhibit multi-layering of cell state markers as described recently^48^, whole-slide analysis shows that there is substantial variation in global tumor organization with some tumors characterized by extensive OPC/NPC differentiation (BT159), extensive AC/MES-like differentiation with minimal OPC/NPC states (BT179), and frequent NPC-like clusters subtly interspersed throughout the tumor (BT145). These findings suggest that multiplexed imaging approaches and spatial analyses will be necessary to fully characterize the global spatial patterning of cell state differentiation in cancers such as GBM which exhibit substantial histologic and genomic heterogeneity across different tumors.

### Primary and micronuclear ruptures are associated with cGAS-STING and interferon signaling

PN and MN ruptures have been associated with DNA damage and innate immune signaling in some settings^18,19^, but the direct downstream consequences of NE rupture have not been characterized in most cancer types, including GBM. While cGAS strongly localizes to PN and MN rupture foci in GBM (**Figure 3)**, whether cGAS leads to activation of downstream cGAS-STING signaling and interferon activation is not well-established *in vivo*, particularly for PN ruptures. Additionally, the DNA damage marker pH2AX is elevated in a subset of PN and MN, with markedly greater signal in MN (**Figure 3**). The cytoplasmic nuclease TREX1 has been proposed to drive NE rupture-associated DNA damage *in vitro* and in breast carcinoma models, but whether it plays a similar role in GBM or other tumors is unknown.

To address these questions, we first stained *LMNB1*-KO U2OS osteosarcoma cell lines, which are known to undergo high levels of spontaneous PN rupture and found that TREX1 localizes to a subset of PN and MN rupture events, as expected (**Figures S7A-S7C**). We then performed t-CyCIF on our glioma TMA (n=145 GBM) using a broad panel of immune, cGAS/STING, DNA damage, and interferon pathway markers as well as antibodies against BAF, lamins, and TREX1 (**Figure 7A**). We observed PN and MN rupture, variable pH2AX signal, and heterogeneous activation of innate immune markers such as pSTAT3, IFITM1/2/3, and MX1 (**Figure 7B**). Exhaustive visual inspection of all cores was required to identify even a small number of PN and MN events with overt co-localization of TREX1 and pH2AX (**Figures 7C** and **7D** and **Figure S7D**). cGAS accumulation was common at BAF-positive PN and MN ruptures (**Figure 7E**), but we were unable to demonstrate STING co-localization to BAF+/cGAS+ nuclear envelopes, despite success in RPE cells grown in culture using the same antibodies (**Figures 2D-2G**). In tissue sections, many tumors cells showed high cytoplasmic STING signal that was difficult to interpret and quantify. However, in GBM cells in which BAF and cGAS co-localized at PN or MN ruptures, we observed evidence of downstream inflammatory activation, including nuclear translocation of IRF3, NF-kB, and pSTAT3, and expression of canonical interferon response proteins such as IFITM1/2/3 (**Figure 7E**). Thus, formation of BAF foci at PN and MN ruptures is very likely associated with activation of the cGAS/STING pathway in at least a subset of cases with consequent activation of transcription factors and genes involved in an interferon response.

**Figure 7:**
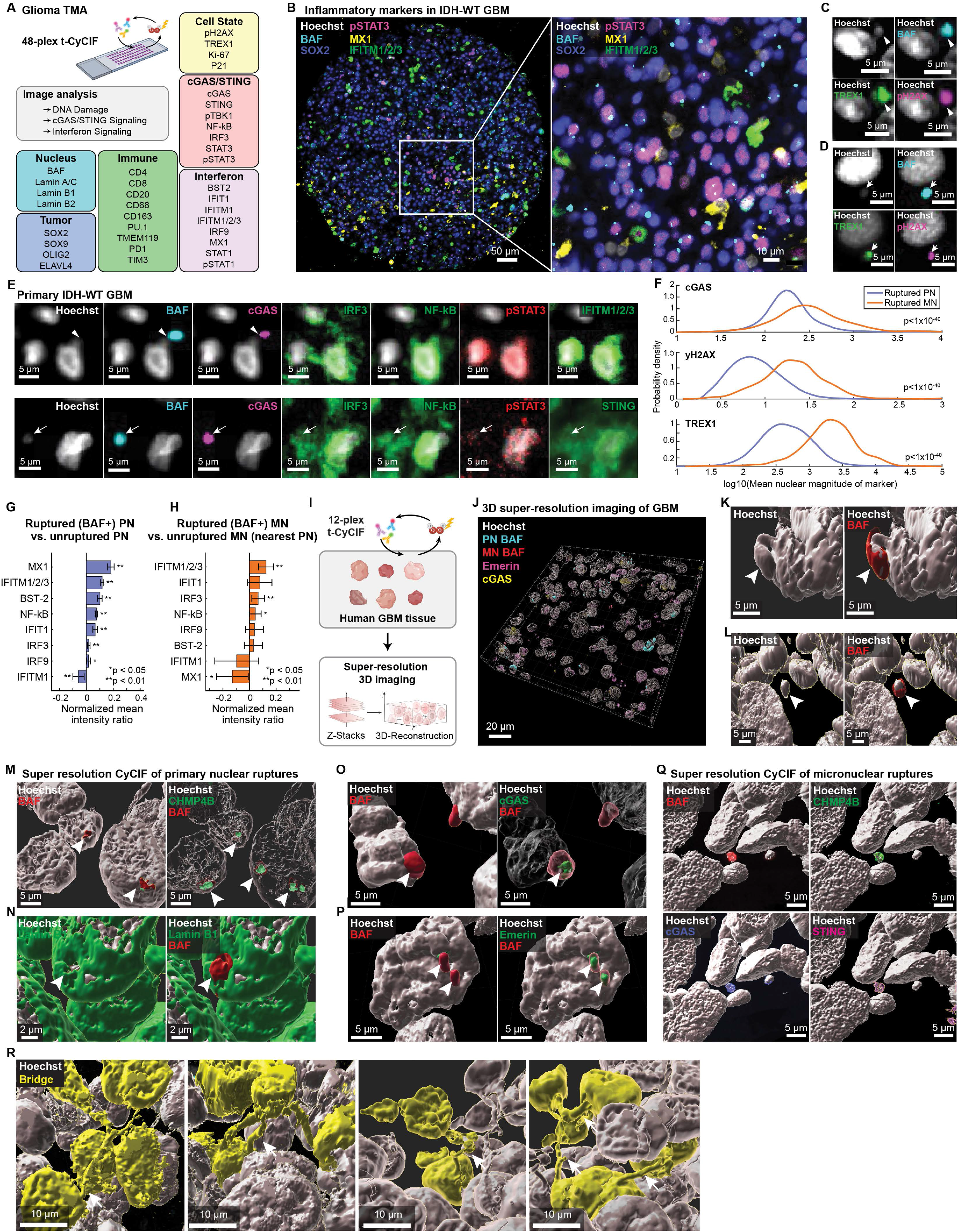
Primary and micronuclear ruptures are associated with cGAS-STING and interferon signaling. **A,** Schematic of t-CyCIF imaging to analyze inflammatory and DNA damage states in the glioma TMA. **B,** Representative CyCIF images of primary IDH-WT GBM showing numerous BAF foci and heterogeneous expression of inflammatory markers in tumor cells. **C,D,** Representative t-CyCIF images showing a large PN bleb (**C**) and MN (**D**) with NE rupture (BAF+) and co-accumulation of TREX1, and pH2AX. **E,** Representative t-CyCIF images of primary IDH-WT GBM cells showing co-localization of BAF and cGAS at ruptured PN (top row) and MN (bottom row), nuclear translocation of IRF3, NF-kB, and pSTAT3, and expression of IFITM1/2/3 (top row). STING showed substantial background staining (bottom row). **F,** Probability density analysis of pH2AX, cGAS, and TREX1 signal in PN and MN BAF foci normalized to Hoechst signal, showing significantly greater signal in MN rupture relative to PN ruptures for each marker (p<1×10^-40^). **G,H,** Quantification of interferon signaling-related markers in 82 primary IDH-WT GBM comparing cells with ruptured and unruptured PN (**G**) and the nearest PN associated with ruptured MN (**H**). p<0.05, **p<0.01; t-test, two-sided, unpaired. **I,** Schematic of super-resolution 3D CyCIF markers and image processing. **J,** Overview of 3D super-resolution CyCIF of 20µm FFPE GBM tissue. **K,L,** Representative 3D surface rendering of primary IDH-WT GBM tumor cells with a BAF-positive ruptured primary nuclear bleb (**K**) and BAF-positive ruptured micronucleus (**L**). BAF forms a ‘shell’ around each ruptured structure. **M-P,** Representative 3D surface rendering of primary IDH-WT GBM tumor cells with PN rupture, showing frequent co-localization of BAF and CHMP4B consistent with ESCRT-III recruitment and repair (**M**). PN ruptures, indicated by BAF, occur at sites of reduced lamin B1 expression (**N**). Super-resolution CyCIF shows frequent co-localization of cGAS (**O**), and Emerin (**P**) at BAF-positive rupture sites. **Q,** Representative 3D surface rendering of primary IDH-WT tumor cells with a micronuclear rupture, showing co-localization of CHMP4B, cGAS, and STING. **R,** Super-resolution CyCIF imaging reveals complex nuclear bridges connecting multiple tumor cell nuclei (arrows; yellow coloration indicates contiguous nuclear structures).

To further analyze the accumulation of DNA damage and cGAS at rupture sites, PN and MN BAF foci were segmented and the normalized intensities of pH2AX and cGAS were quantified relative to DNA content. We found that pH2AX, cGAS, and TREX1 showed a continuous distribution of signal at BAF+ PN and MN rupture foci, suggesting varying levels of activation. Further analysis showed that MN ruptures exhibited substantially higher mean pH2AX (3.7-fold greater; p=1.1×10^-6^), cGAS (6.8-fold greater; p=2.2×10^-17^), and TREX1 (5.7-fold; p<1×10^-40^) than PN ruptures normalized to DNA content (**Figure 7F**). Consistent with robust accumulation of cGAS, analysis of PN (**Figure 7G**) and MN (**Figure 7H**) showed that both types of rupture were associated with NF-kB and IRF3 nuclear translocation and increased IFITM1/2/3 expression. Thus, as with prior experiments (**Figure 3**), while pH2AX and cGAS may accumulate at both PN and MN ruptures, MN are associated with substantially greater levels of DNA damage, TREX1 localization, and cGAS-STING activation, and we show the first evidence in human tissues that cGAS-STING activation likely leads to downstream interferon signaling through canonical effectors such as NF-kB and IRF3.

To examine the relationship between ruptures and subcellular localization of effectors such as cGAS and STING, the ESCRT-III complex, and nuclear lamins, we performed 3D CyCIF using a 12-marker panel on a Zeiss LSM 980 confocal microscope with Airyscan2 imaging in super-resolution mode on 4 specimens (**Figure 7I** and **Figures S7E** and **S7F**). Fine mapping of BAF-positive rupture events showed that BAF formed a tight ‘shell’ around sites of ruptured PN, often occurring at regions of high curvature. These occasionally corresponded to large blebs in the PN envelope with substantial protrusion (**Figures 7J** and **7K**). Tight shells of BAF were also observed around ruptured micronuclei which typically comprised a compact collection of chromatin (**Figure 7L**). Nearly all BAF foci were associated with focal accumulation of CHMP4B, a component of the ESCRT-III repair complex, which is involved in NE repair (**Figure 7M**). Evaluation of nuclear lamins revealed irregular shells of lamin B1, with PN rupture events occurring predominantly in multifocal regions exhibiting reduced lamin B1 protein (**Figure 7N**). While cGAS accumulation was observed in only a subset of ruptures by conventional widefield t-CyCIF data (**Figure 3**) super-resolution imaging showed that nearly all BAF foci exhibited at least subtle cGAS accumulation (**Figure 7O**). Most ruptures exhibited BAF co-localization with emerin (particularly at PN ruptures), confirming that BAF likely recruits additional LEM-domain binding proteins of the A-type lamina complex to rupture sites (**Figure 7P**).

We also observed local accumulation of CHMP4B and cGAS at nearly all BAF-positive MN, as well as focal accumulation of STING, further suggesting that widefield imaging inherently under-samples these phenotypes relative to high-resolution 3D imaging (**Figure 7Q**). However, despite the increased resolution of Airyscan2 imaging, we did not observe STING accumulation at BAF-positive PN ruptures – not even those with cGAS accumulation. Thus, it is possible that STING does not focally accumulate at PN ruptures or that activation of downstream interferon signaling (**Figures 7E** and **7G**) may be driven by other mechanisms in these cells. Fine mapping of chromosomal bridges in these specimens also revealed the presence of structures that have not been widely studied, including nuclei with multiple bridging structures, and ‘syncytial’ collections of >2 nuclei connected by multiple DNA bridges, indicating that dividing glioma cells may undergo additional complex events that impact chromatin integrity which will require further classification (**Figure 7R**).

## Discussion

A fundamental challenge in studying the mechanisms of human disease is that cellular processes are best understood in cultured cells and animal models that can readily be subjected to genetic and drug perturbation followed by live-cell analysis. However, the chemical and mechanical environment of human tissues and tumors is radically different from that of cells in culture or even mouse models. For instance, rates of cell division are highly discordant between cultured cells and tumor tissues.^23,49^ This leads to significant uncertainty in translating - from a dish to a tissue - knowledge of even fundamental and highly studied mechanisms such as cell division and chromosome segregation. The current work builds a bridge between data and models of NE rupture *in vitro* and the nuclear atypia observed in tissues by combining live-cell and multiplexed immunofluorescence of cell lines and organoids with classical histology and high-plex tissue profiling of human tumor specimens. Multiplexed measurement is particularly important in this setting both as a means to deconvolve the complex multicellular environment of actual tumors, and as means of inter-relating fixed human tissues and *in vitro* cell line-based measurements. The ability to combine CyCIF with high-resolution confocal microscopy has also been critical because many of the processes involved in NE rupture involve small structures and highly localized protein accumulation (e.g., BAF and cGAS foci). 3D imaging of sections that are thick enough to encompass whole cells further enables detection of phenotypes such as chromosome bridge formation that are difficult or impossible to characterize in conventional thin tissue sections. In addition, analysis of combinations of protein markers provides a means of linking cell states identified in images to those defined by scRNA-seq signatures (and *vice versa*). The bridges we build between *in vitro* and *in vivo* studies, and between RNA and immunofluorescence-based cell-state definitions make it possible to dissect the intricacies of nuclear envelop biology in the intermixed and heterogeneous environments of the human tumor microenvironment.

We find that NE rupture events, including rupture of the primary nuclear (PN) and micronuclear (MN) membranes are common in GBM and several other types of cancer: 10-25% of cells show evidence of PN rupture and 1-10% of MN rupture. We also provide direct evidence, in a native tumor context, of chromosomal bridges that recruit cGAS and induce innate immune signaling responsible for patterning the TME. The frequency of nuclear envelope rupture suggests that inhibition of the machinery that detects and repairs damage to the NE, BAF and the ESCRT-III complex for example, may lead to catastrophic loss of nuclear integrity and cell death in tumors with high degrees of instability, making such machinery an attractive target for novel therapeutic agents.^50^

GBMs comprise a mixture of different cell types including cells with the properties of oligodendrocyte progenitors (OPC), neural progenitor (NPC), astrocyte (AC), and mesenchymal cells (MES).^46^ The properties of these lineages have been most thoroughly described using dissociative single-cell RNA sequencing. However, acute NE rupture does not have a defined transcriptomic signature and scRNA-seq cannot detect the morphological changes that best define the rupture process. Moreover, even with advances in spatial transcriptomics, single-cell transcriptional analysis of archival human tissue remains challenging and its ability to detect structural phenotypes or changes in localization of effectors is limited.^51^ We therefore used scRNA-seq data to identify candidate genes that are differentially expressed in GBM lineages and develop a multiplexed antibody panel to label their protein products. We tested the panel using TMAs and tumors generated with patient-derived cell lines whose composition had been established by RNA-Seq. In addition, we validated antibodies for different lamins and NE proteins using cultured cells that had been subjected to a wide range of perturbations designed to alter NE function and structure. This yielded several cross-validated 12 to 45-plex Ab panels that effectively discriminated cell states and NE proteins in both cell lines and histological preparations. Using partly redundant and overlapping markers of cell states and subcellular processes reduces the likelihood of errant classification due to a marker that is either not detected, protein expression that is absent due to genetic alteration or variable biology, or sub-populations of cells with distinctive or hybrid phenotypes, such as EGFR^high^ NPC-like cells.

This work revealed that NE rupture is strongly associated with reduced levels of nuclear lamins in PN and MN, and that nuclear lamin levels correspond with the underlying differentiation state of GBM tumor cells. MN and PN ruptures are most common in NPC-like cells. PN rupture is associated with low lamin A/C expression globally throughout the nuclear envelope and with low lamin B1 focally at the specific point of rupture, while MN rupture is associated with reduced lamin B1 and B2 in the micronuclear envelope and increased cell proliferation. PN and MN ruptures have been reported to lead to innate immune activation.^14,52^ In tumor tissues, we observe that cells with ruptured PN and MN have robust accumulation of cGAS at BAF foci. We further demonstrate increased nuclear translocation of IRF3, NF-kB, and pSTAT3, and expression of interferon response genes (proteins) such as IFITM1/2/3 associated with PN and MN ruptures in human tumors. In tissues, cGAS was approximately six-fold more abundant in ruptured MN than PN and STING accumulation was only observed at MN ruptures. Biochemical studies show that cGAS activation is transiently inhibited by chromatin^53^ potentially explaining the longer duration of MN ruptures relative to PN ruptures (**Figure S1**), and the consequent increase in exposure of MN chromatin to the cytoplasmic compartment where it would induce STING localization and activation. In our studies, super-resolution 3D CyCIF was necessary to quantify the relationship among focal reductions in B-type lamina components such as lamin B1, B2, and nucleoporins (NUP133), STING localization, CHMP4B and cGAS localization, and morphologic analysis of atypical and ruptured PN and MN. Thus, this work provides an example of how multi-scale analyses using both 2D and 3D methods generate complementary information and a more comprehensive assessment than either method alone.

The causes and consequences of NE rupture in cancer have been primarily studied in established cell culture models (e.g., RPE, U2OS, HeLa cells)^17^ using live-cell imaging and microfluidic confinement and in xenografts (breast carcinoma and melanoma).^18^ These studies have arrived at conclusions broadly congruent with ours. However, tissue studies show some discrepancies with respect to the link between PN rupture and DNA damage. We find that DNA damage (pH2AX staining) is associated, in RPE cells, with a substantial number of MN ruptures but not PN ruptures, except in the case of XR exposure, which show increased damage overall, likely directly related to XR-treatment. However, in untreated primary human GBM tissues, pH2AX accumulation is observed in association with both PN and MN ruptures, albeit with substantially higher levels in MN. Such differences may reflect variation in DNA damage repair *in vivo* and *in vitro* or incidental association of DNA damage and PN rupture in rare cells *in vivo* given the much slower division rate than in cultured cells. Of note, segmentation and analysis of a large number of rupture events in human tissue demonstrates pH2AX staining over a wide and continuous range of signal intensities. Imposition of a dichotomous score on a continually varying level of DNA damage, without standardized thresholds for ‘positive’ signal (which may vary by tissue and context) is almost certain to lead to substantial variability between datasets and interpretation of damage incidence. We are nonetheless confident, from analysis of BAF-positive PN and MN ruptures in tissues using multiple imaging modalities, including super-resolution imaging in which the details of the NE are visible, that MN exhibit substantially higher levels of pH2AX signal than PN ruptures, particularly in BAF+ ruptured MN. This strongly suggests that MN ruptures are more susceptible to DNA damage than PN ruptures in GBM tissues. Previous studies of breast carcinoma tissues and cell line models^19^ reported frequent accumulation of the TREX1 nuclease at PN and MN rupture sites and hypothesized that this drove the DNA damage associated with NE rupture. We find that TREX1 signal may be identified with a continuous distribution in PN and MN rupture events, with substantially greater signal in MN, though overt focus formation was only rarely observed. Such differences may result from reagents, criteria used to gate marker intensities in imaging data, and other experimental and biological variables.

Using antibody-based lineage markers, we find that GBM tumors exhibit both short and long-range patterning of lineage cell states^26^ that appears to reflect the dominant lineage program present in the tumor cells themselves. For example, tumors that have robust expression of AC-like markers can exhibit multi-layered patterning in which MES-like cells surround areas of microvascular proliferation and regions of necrosis while NPC/OPC cells preferentially localize at infiltrative edges (the interface between the tumor and the brain). However, tumors with NPC/OPC-predominant expression exhibit diffuse expression of NPC and OPC markers throughout the tumor mass, scant necrosis, and a paucity of AC and MES-like cells. Accordingly, while some spatial relationships between cell state and anatomic organization appear to be consistent across tumors, such as the association of MES-like cells with peri-necrotic regions, the global patterning of tumors is tied to the underlying cell state differentiation of tumor cells. This patterning of tumor lineage corresponds to the pattern of NE instability, with greater levels of NE rupture in NPC-like regions, which are associated with lower lamin A/C expression and higher proliferation.

Collectively, our data show that NE rupture is surprisingly common in a wide variety of human tumors, and strongly tied to nuclear lamin protein expression, lineage and cell state, and activation of innate immune signaling. Such events may condition the tumor immune microenvironment to support tumorigenesis, and lead to DNA damage and chromosomal alterations that drive tumor progression. Given the high rate of NE instability observed in many tumors and cell lines, targeted disruption of NE repair processes may represent a potential therapeutic opportunity.

### Limitations of the Study

Our study provides the first large-scale characterization (“Atlas”) of nuclear envelope rupture rates in human cancer tissues from hundreds of specimens and validates BAF focus formation as a biomarker of PN and MN rupture. However, imaging of histological preparations represents only a single point in time and correlation of fixed and live-cell imaging of cultured cells shows that fixed cell data underrepresent the true frequency of NE rupture due to ongoing repair. *Ex vivo* tissue slice experiments may be necessary to track the incidence of nuclear instability and provide additional context on drivers of NE rupture, such as mechanical deformation of cells which is difficult to infer from tumor specimens. The lack of a detailed temporal description of the coordinated molecular fluctuations that occur during NE rupture and repair limits the use of ergodic principles which enable reconstruction of dynamic processes from fixed-time data of tissues.^23^

## Acknowledgements

This work was supported by the Ludwig Cancer Research and the Ludwig Center at Harvard (P.K.S., D.P, S.S.) and by NCI grants U54-CA225088 and U2C-CA233262 (P.K.S., S.S.). Development of CyCIF imaging methods and image processing software is supported by a Team Science Grant from the Gray Foundation (P.K.S., S.S.), the Gates Foundation grant INV-027106 (P.K.S.), the David Liposarcoma Research Initiative (P.K.S., S.S.), Emerson Collective (P.K.S.). S.C. is supported by training grants T32-GM007748 (NIGMS) and T32-CA009216 (NCI). S.S. is supported by the BWH President’s Scholars Award. D.P. is supported by HHMI. KLL is supported by NCI grants P50-CA165962, R01-CA188228, R01-CA219943, U19-CA264504, 3000 Miles to the Cure, and the National Brain Tumor Society. We thank Shu-Hsien Sheu, Robert Krueger, Donglai Wei, and Terri Woo for their expert assistance.

## Author Contributions

SC, BC, JSL, RR, LB, YX, AS, and CY performed experiments and imaging.

SC, BC, JSL, RR, LB, SSC, CY, GJB, SJC, and ES performed data analysis.

SC, SS, and PKS wrote the paper and all authors reviewed drafts and the final manuscript.

SC, JBT, and SS prepared the figures.

SS and KLL supervised clinical research.

SS, PKS, KLL, AS, and DP supervised the overall research.

## SUPPLEMENTAL FIGURE TITLES AND LEGENDS

**Figure S1: Characterization of primary and micronuclear ruptures across human cancers and cell lines. A,** Representative BAF immunohistochemistry (IHC) images from human TMA for high-grade serous (ovarian) carcinoma (HGSC), breast carcinoma (BRCA), lung adenocarcinoma (LUAD), diffuse large B-cell lymphoma (DLBCL), pancreatic ductal adenocarcinoma (PDAC), prostate adenocarcinoma (PRAD), colorectal adenocarcinoma (COAD), and melanoma (MEL). **B,** Representative image of BAF immunofluorescence (IF) in a primary IDH-WT GBM, showing numerous BAF foci indicative of PN and MN rupture. **C,D** Hoechst staining of PN (**E**) and MN (**F**) rupture examples from Figure 1 demonstrating nuclear morphology. **E,** Representative 3D confocal imaging of primary IDH-WT GBM tissue, showing BAF rupture foci (cyan) at sites of reduced lamin B1 expression. **F,** Representative 3D expansion microscopy IF imaging of primary IDH-WT GBM tissue, showing PN BAF foci (cyan) at sites of reduced lamin B1 protein and chromatin herniation consistent with blebs. **G,** Schematic representation of the FIB-SEM microscope used to prepare HGSC sample by milling trenches to isolate a 35×12×27 um volume and mill the imaging area every 20 nm along the z plane to capture 1309 serial scanning electron images. **H,** Representative time-lapse imaging of a BAF-eGFP GBM PDCL (BT179) stained with silicon rhodamine DNA dye (SiR-DNA) showing PN distortion during migration, blebbing of the NE, and formation of a BAF focus with focal leakage of SiR-DNA dye (arrowheads). **I,** Representative time-lapse imaging of BAF-eGFP transgenic PDCL (BT145, BT159, BT179 and Kuramochi cells stained with SiR-DNA undergoing mitosis formation and rupture of MN with formation of intense BAF foci. **J,** Representative time-lapse imaging of BAF-eGFP transgenic BT159 and Kuramochi cells stained with SiR-DNA undergoing mitosis with chromosomal mis-segregation and formation of a chromosomal bridges. **K,L,** Phase-contrast/immunofluorescence 2D imaging of BAF-eGFP transgenic Kuramochi ovarian carcinoma cells (**K**) and segmentation (**L**) showing multiple time-points of a MN rupture event. **M-N,** Time-course intensity plots of BAF-eGFP signal in segmented PN (**M**) and MN (**N**) rupture events in Kuramochi cells (total imaging time=18 hours). These data show that both event types tend to resolve with loss of BAF signal, and PN rupture events (mean length 85 minutes, n=6) are typically substantially shorter than MN rupture events (mean length 253 minutes, n=25) (p=0.014) (t-test, two-sided, unpaired). **O,P,** Comparison of rupture events by BAF area (in pixels) shows that larger rupture events persist significantly longer than shorter events (linear regression; p=0.003, R=0.5153). **Q,R,** Comparison of rupture events by mean BAF signal intensity in the segmented area shows that more intense rupture events persist significantly longer (linear regression, p=0.0467, R=0.3599).

**Figure S2: Pan-cancer expression and dependency of lamin genes in cancer. A-C,** TCGA Pan-*Cancer* data (n=10,071 tumors, 30 subtypes) show that GBM and gliomas have low expression of *LMNA* mRNA compared to other subtypes (49.5% of mean *LMNA* expression compared to the mean of all tumors; p=1.25×10^-56^), but moderate levels of *LMNB1* (0.28 log2 fold change; p=2.90×10^-4^) (**B**) and *LMNB2* (−0.21 log2 fold change; p=1.20×10^-3^) (**C**) (t-test, two-sided, unpaired). **D,E,** Mutation (**D**) and copy-number analysis (**E**) of TCGA Pan-Cancer data (n=10,071 tumors, 30 subtypes) show only very rare alterations at the *LMNA* gene locus in cancer. **F,G,** Analysis of DepMap gene dependency studies in CCLE lines (n=1095) show that GBM and other gliomas are among the most dependent on *LMNA* expression for cell viability in CRISPR (p=1.43×10^-20^; n=49) (**F**) and RNAi (p=3.43×10^-11^; n=31) (**G**) experiments relative to the mean of all cell lines (t-test, two-sided, unpaired). **H,I,** Glioma and soft tissue lines show some dependency on *LMNB1* expression in CRISPR (p=4.06×10^-6^; n=68) (**H**), but not RNAi (**i**) experiments (t-test, two-sided, unpaired). **J,K**, There was no strong dependence on *LMNB2* in glioma cell lines, though melanoma lines show a slight dependence. **L,** Data from the glioma TMA t-CyCIF (Fig. 3), were used to compare primary IDH-WT GBM (n=82) vs. recurrent/residual IDH-WT GBM (n=45), showing that recurrent cases exhibit significantly higher levels of lamin A/C (p<0.0001). **M,** Comparison of all primary and recurrent IDH-WT GBM (n=127), and all primary and recurrent IDH-mutant GBM (n=15) in the glioma t-CyCIF TMA data show that IDH-mutant cases exhibit significantly lower levels of lamin A/C (p<0.0001) and significantly higher PN rupture rates (p<0.0001) compared to IDH-WT GBM (t-tests, two-sided, unpaired).

**Figure S3: Classification of glioblastoma cell states by single-cell RNA-sequencing.** Single-cell RNA-seq data from 28 adult and pediatric GBM (Neftel *et al.*) were classified according to lineage and cell state, including AC-like, MES-like, OPC-like, and NPC-like. These data were evaluated for cell state-specific markers that may be amenable to t-CyCIF. **A-D,** Two-dimensional scRNA-seq cell state graphs for candidate AC-like (**A**), MES-like (**B**), NPC-like (**C**), and OPC-like (**D**) markers, showing significant enrichment in each cell state (t-tests, two-sided, unpaired). **E,** scRNA-seq plot of SOX2, which was expressed by all lineages and used as a general marker of tumor cells in t-CyCIF (t-test, two-sided, unpaired).

**Figure S4: Spatial analysis of nuclear instability and cell state patterning in GBM autopsy specimens. A,** t-CyCIF images of MES-like markers HIF1A and NDRG1 in peri-necrotic cells in IDH-WT GBM autopsy tissue. **B,** Representative t-CyCIF imaging of four human autopsy tissue specimens, showing the patterning of cell state markers SOX10 (OPC-like), ELAVL4 (NPC-like), HOPX (AC-like), and NDRG1 (MES-like). **C,** Spatial analysis of marker expression in human autopsy specimens. Graphs extend from the annotated tumor center (left-hand side) to the tumor edge (line), to adjacent normal tissue in concentric circles. **D,** Transcriptomic data from the IVY Glioblastoma Atlas Project (IVY-GAP). Heatmap shows microregional spatial transcriptomic data of lamin genes and GBM cell states from seven anatomically distinct tumor regions (n=10 GBM).

**Figure S5: Spatial analysis of nuclear instability and cell state patterning in GBM PDX. A,** Representative low-, medium-, and high-power t-CyCIF imaging of coronal brain sections from a patient-derived xenograft model generated from the BT112 PDCL line. SOX2 is used as a pan-tumor cell marker showing the hemispheric anatomic distribution of the tumor with extension into the corpus callosum. t-CyCIF shows reduced lamin A/C at the tumor periphery, corresponding with increased BAF rupture and NPC-like differentiation. High-power views demonstrate a cluster of NPC-like differentiation showing low lamin A/C, increased PN rupture (BAF foci), NPC-like markers (ELAVL4+, DCX+), and surrounding cells from other lineages. **B,** Representative low-, medium-, and high-power t-CyCIF imaging of coronal brain sections from a patient-derived xenograft model generated from the BT145 PDCL line. SOX2 is used as a pan-tumor cell marker showing the bi-hemispheric anatomic distribution of the tumor which crosses the midline. Medium-power views from the BT145 PDX tumor show numerous intermixed clusters of NPC-like cells (DCX+/ELAVL4+/INSM1+), surrounded by AC-like cells (GFAP+/EGFR+/SOX9+). High-power views of NPC-like tumor cell clusters show that they exhibit low lamin A/C expression and increased PN rupture (BAF+ foci) relative to surrounding AC-like and OPC-like cells.

**Figure S6: Additional spatial analysis of nuclear instability and cell state patterning in GBM PDX. A,** Representative low-power t-CyCIF imaging of coronal brain sections from a patient-derived xenograft model generated from the BT159 PDCL line. SOX2 is used as a pan-tumor cell marker showing the midline anatomic distribution of the tumor. Scalebars, 500µm. Medium-power views from the BT159 PDX tumor show diffuse OPC-like and NPC-like differentiation with many cells exhibiting markers from both cell states. Scattered AC-like cells (SOX9+) are present in the background. High-power views of tumor cells show that those with NPC/OPC-like expression exhibit low lamin A/C expression, PN ruptures, and expression of associated cell state markers (SOX10, OLIG2, DCX). **B,** Representative low-power t-CyCIF imaging of coronal brain sections from a patient-derived xenograft model generated from the BT179 PDCL line. SOX2 is used as a pan-tumor cell marker showing the midline anatomic distribution of the tumor which invades the brainstem and cerebellar white matter with a diffusely infiltrative growth pattern. High-power views of tumor cells show predominant expression of AC-like markers (SOX9, GFAP, EGFR) and MES-like markers (NDRG1) with limited expression of NPC and OPC-like markers by tumor cells.

**Figure S7: Nuclear instability is associated with DNA damage, innate immune activation, and complex nuclear structure in glioblastoma. A,** *LMNB1*-ko U2OS (osteosarcoma) cells immunolabeled with TREX (green) and BAF (red) antibodies. **B,C,** Immunofluorescence data show that BAF+ foci occasionally exhibit co-localization of TREX1 in PN (B) and MN (**C**). **D,** Representative Hoechst imaging demonstrating the morphology of a large PN bleb (from Fig. 7C) which shows co-accumulation of BAF, TREX, and pH2AX in IDH-WT GBM. **E,** Representative Z-slices from 3D super-resolution CyCIF reconstruction of a region of IDH-WT GBM, showing nuclear morphology, BAF+ rupture events, and other phenotypes. **F,** Representative 3D super-resolution CyCIF showing combinations of markers in reconstructed 20µm IDH-WT GBM tissue from four resection specimens.

## METHODS

### RESOURCE AVAILABILITY

#### Lead contact

Further information and requests for resources and reagents should be directed to and will be fulfilled by the lead contact, Sandro Santagata MD, PhD.

#### Materials availability

This manuscript contains no unique reagents or resources; all antibodies are available commercially (see Table S3 and key resources table).

#### Data and code availability

All images and derived data will be accessible without restriction through the National Cancer Institute Human Tumor Atlas Network (HTAN) Portal (https://humantumoratlas.org/). The HTAN participant (specimen) ID numbers will be found in **Table S1**. Additionally, other data from this study will be available through an index page on GitHub, which will be archived on Zenodo at the time of acceptance.

- Image and data analyses were performed using custom scripts in Python, ImageJ, and MATLAB. The entire codebase is accessible under an MIT open-source license through an index page on GitHub, which will be archived on Zenodo (DOI) at the time of acceptance.
- All data necessary to reanalyze the data should be available via Zenodo or HTAN. For issues accessing the data reported in this paper, please contact the Lead Contact.

## EXPERIMENTAL MODEL AND STUDY PARTICIPANT DETAILS

### Human participants

Our research complies with all relevant ethical regulations and was reviewed and approved by the Institutional Review Boards (IRB) at Brigham and Women’s Hospital (BWH), Harvard Medical School (HMS), and Dana Farber Cancer Institute (DFCI). Discarded human formalin fixed paraffin embedded (FFPE) tissue samples were used after diagnosis under excess tissue discarded tissue protocol 2018P001627 (reviewed and managed by the Mass General Brigham Institutional Review Board) which waives the requirement for patient consent. Genomic analysis of adult glioblastoma tissues by OncoPanel sequencing was performed after written informed patient consent had been obtained under DFCI IRB protocol 10-417 or 17-000. Adult glioblastoma tissue samples used to generate patient derived xenografts and cell lines by the DFCI Center for Patient Derived Models (CPDM) were obtained following written consent to DFCI IRB protocol 10-417 or 17-000. The study is compliant with all relevant ethical regulations regarding research involving human tissue specimens. The Principal Investigator is responsible for ensuring that this project was conducted in compliance with all applicable federal, state, and local laws and regulations, institutional policies, and requirements of the IRB.

Formalin fixed and paraffin embedded (FFPE) tissue specimens were retrieved from the archives of the Department of Pathology of Brigham and Women’s Hospital with Institutional Review Board (IRB) approval under DF/HCC Protocol #10-417 or waiver of consent protocols. Commercial human tissue microarrays (TMAs) were purchased from Pantomics. All glioma cases were reviewed and classified according to the revised W.H.O. 2021 Classification of Tumours of the Central Nervous System (S.C., S.S.). The characteristics of cases including demographics, genotyping, immunohistochemical, and clinical data are provided in **Table S1** in accordance, when possible, with emerging metadata standards for tissue imaging^54^ and IRB guidelines. For glioblastoma, specimens imaged by CyCIF included 145 GBM tissues, including 127 IDH-WT GBM (W.H.O. grade 4) (82 primary, 45 recurrent) and 15 IDH-mutant GBM (also known as Astrocytoma, IDH-mutant, W.H.O. grade 4) (4 primary, 11 recurrent). The TMA also included 14 oligodendroglioma, IDH-mutant, 1p/19q co-deleted specimens (W.H.O. grade 2) (12 primary and 2 recurrent tumors); 15 anaplastic oligodendroglioma, IDH-mutant, 1p/19q co-deleted specimens (W.H.O. grade 3) (6 primary and 9 recurrent tumors). Four additional glioblastomas obtained from brain autopsies were used for whole slide imaging. For other tumors, commercial tissue microarrays were purchased from Pantomics (https://www.pantomics.com/), including pancreatic cancer (PAC1021), breast carcinoma (BRC2281), lung carcinoma (LUC2281), prostate carcinoma (PRC1021), melanoma (MEL961), lymphoma (LYM1021). For colorectal carcinoma, we stained an in-house tissue microarray (HTM402) containing colorectal adenocarcinoma specimens. For ovarian carcinoma, we stained a tissue microarray generated by the Helsinki University (405.2)

### Patient-derived Cell Line and Xenografts

Patient-derived cell lines (PDCL) and patient-derived xenograft (PDX) tissues of glioblastoma for BT112, BT145, BT159, and BT179 and related mRNA expression profiling data were obtained from the DFCI Center for Patient Derived Models (CPDM) which generates, characterizes, stores, and distributes cancer models.

## METHOD DETAILS

### Immunohistochemistry

Tissue sections were deparaffinized and rehydrated through xylene and graded alcohols. Endogenous peroxidase activity was blocked by incubating sections for 15 minutes in 3% hydrogen peroxide and 100% alcohol (1:1). Heat induced antigen retrieval was carried out with Dako citrate buffer (pH=6.0) and pressure cooker at 122 +/-2°C for 45 seconds (15+/-2 PSI). Sections were incubated at room temperature for 45 minutes with BANF1 antibody (Abcam, EPR7668, rabbit monoclonal) at 1:4000 dilution. Application of the primary antibodies was followed by incubation for 30 minutes with Dako Labeled Polymer-HRP anti-rabbit IgG as a secondary antibody (K4011), and visualized with 3, 3’– diaminobenzidine (DAB) as a chromogen (Envision+ System) and hematoxylin counterstaining. Primary nuclear (PN) and micronuclear (MN) BAF foci in human tissue and cell line specimens were manually quantified by a pathologist (S.C.) in human tissue sections via conventional light microscopy (Olympus CX41 microscope).

### RPE Cell Line Growth and Perturbations

Human retinal pigmented epithelium (hTERT RPE-1; ATCC CRL-4000) cells grown in RPMI medium with GlutaMAX supplement with 10% fetal bovine serum and 1% penicillin/streptomycin. Cells were perturbed with each of the following conditions: BAF siRNA transfection, LMNB1 siRNA transfection, HT-DNA transfection, MPS1 inhibitor (NMS-P715, 1μM), and X-radiation (XR, 2Gy).

### Lentivirus transduction

GBM PDCL (BT145, BT159, BT179) and Kuramochi HGSC cell lines were tranduced with BAF-eGFP (Addgene # 101772) lentivirus, using Viraductin reagent (Cellbiolabs, AAV-200). Transduced cells underwent fluorescence-activated cell sorting (FACS), into quartiles by nuclear BAF expression. The bottom quartile of BAF-eGFP positive cells were selected and expanded to minimize transgene-induced NE instability, with visual confirmation of nuclear expression and focus formation. A subset of BAF-eGFP lines (BT145, BT159, BT179) underwent transduction with NLS-RFP lentivirus using Viraductin reagent (Cellbiolabs, AAV-200) to generate BAF-eGFP / NLS-RFP dual reporter lines.

### PDCL Cell Culture

BT112, BT145, BT159, and BT179 glioblastoma cell lines with BAF-GFP reporter were cultured non-adherently at 37°C and 5% CO_2_ in Corning® Ultra-Low Attachment 75cm² U-Flasks (Corning, Catalog #3814). Media consisted of NeuroCult™ NS-A Proliferation Kit (StemCell, Catalog #05751) supplemented with 0.1% Heparin (StemCell, Catalog #07980), 0.01% bFGF (StemCell, Catalog #78003) and 0.02% EGF (StemCell, Catalog #78006). Media was changed every 3 days and cells were passaged approximately every week.

### PDCL Live-Cell Imaging

96-well imaging plates (Ibidi, Catalog #89626) were coated with R&D Systems™ Cultrex™ Mouse Laminin I Pathclear™ (Fisher Scientific, Catalog #34-000-1002) to ensure a monolayer culture and incubated at 37°C overnight. Cells were plated at 100k cells/cm² for all cell lines for quantification of rupture frequency. For SiR-DNA experiments, SiR-DNA was added to the well media at a 100nM concentration. After incubating 24 hours, cells were imaged on a GE IN Cell Analyzer 6000 at 2–3-minute time lapse intervals at 20X magnification (NA 0.75) for 24 hrs. BAF-GFP was visualized using Blue (488 nm) excitation and standard FITC emission filters (524 nm), and SiR-DNA with Red (642 nm) excitation and standard Cy5 emission filter (682 nm). Temperature was maintained at 37°C for the duration of imaging. Time lapse images were compiled utilizing a custom MATLAB script. Images were viewed and annotated manually via visual inspection in Fiji.

### Tissue-Based CyCIF Acquisition Protocols

FFPE sections of GBM PDX samples were prepared and stained with a 36-plex antibody panel according to the previously described CyCIF protocols^55^ (see **Table S3**).

### Baking and dewaxing

To prepare samples for antibody staining, slides were automatically baked at 60°C for 30 min, dewaxed at 72°C in BOND Dewax Solution, and antigen retrieval was performed at 100°C for 20 min in BOND Epitope Retrieval Solution 1 (ER1) by the Leica BondRX machine.

### Photo-bleaching and autofluorescence reduction

After baking and dewaxing, the slides were immersed in a bleaching solution (4.5% H2O2, 20 mM NaOH in PBS) with LED light exposure for 1 hr to reduce autofluorescence. To mitigate non-specific antibody binding, slides were washed for 6 times with 1X PBS for 10-15s and then incubated overnight with secondary antibodies (anti-mouse, and anti-rabbit, see **Table S3**) diluted in 2 mL of SuperBlock™ Blocking Buffer (Dilution 1:1000; ThermoFisher Scientific, cat#37515) at 4°C in the dark. Slides were subsequently washed 6 times with 1X PBS before photobleaching them again for 1 hr.

### Antibody staining, slide mounting, and imaging

For each CyCIF cycle, samples were incubated overnight at 4°C in the dark with Hoechst 33342 (1:10,000; ThermoFisher Scientific, cat# 62249) for nuclear staining along with either primary conjugated antibodies or primary unconjugated antibodies (see **Table S3** for antibody information and dilutions) in 2mL of SuperBlock™ Blocking Buffer (ThermoFisher Scientific, cat#37515). Incubation with primary unconjugated antibodies was followed by secondary antibody incubation at room temperature for 1 hr in the dark. For CyCIF antibodies that were only available from vendors as primary unconjugated antibodies, custom conjugates were requested from Cell Signaling Technology. After staining, slides were washed for 6 x 10 seconds, mounted with 24 x 60 mm coverslips using 150 µL of 50% glycerol, and then dried at room temperature for 30 minutes. Once coverslipped, slides were automatically imaged using a RareCyte Cytefinder II HT using the following channels: UV, cy3, cy5, and cy7 (these are nominal names since Cy3/5/7 dyes are not used in CyCIF; see **Table S3** for actual fluorophores). Imaging was performed with the following parameters: Binning: 1 x 1; Objective: 20X; Numerical Aperture: 0.75; Resolution: 0.325 mm/pixel. Image exposures were optimized for each channel to avoid signal saturation but kept constant across samples. To demount coverslips between CyCIF cycles, slides were immersed in containers of 1X PBS (5 slides per container) for 10 min. Before the subsequent cycle of antibody staining, slides were photobleached for 1 hr as described to deactivate the fluorophores and washed 6 x 15 seconds in 1X PBS to wash off the bleaching solution. Detailed protocol of CyCIF is available in protocols.io (https://doi.org/10.17504/protocols.io.bjiukkew).^56^

### Plate-based CyCIF imaging

RPE cells were plated into 96-well glass bottom plates (3 wells per condition) and plate-based CyCIF was performed as described in Lin et al., 2016.^57^ Cells were fixed with 4% PFA at room temperature (RT) for 30 minutes, permeabilized with ice-cold methanol at RT for 10 min, washed with PBS three times, and blocked with 50 µL of Odyssey blocking buffer at RT for 1 hr. Six rounds of antibody incubation, imaging, and fluorophore inactivation were then performed. All antibodies were diluted 1:100 and incubated overnight at 4°C in the dark followed by staining with Hoechst 33342 for 10 min at RT. For the first cycle of imaging, secondary antibodies were diluted 1:1000 and incubated for 1 hr at RT. See **Table S3** for a complete list of all antibodies used. Fluorophore inactivation was performed by incubating cells with a PBS solution with 4.5% H2O2 and 20 mM NaOH for 1 hr under an LED light. Images were acquired with the GE IN Cell Analyzer 6000 using an 40X/0.95 NA objective lens and the following filter set: ‘DAPI channel’ with 455-nm peak excitation/25-nm half-bandwidth, ‘488 channel’ with 525-nm peak excitation/10-nm half-bandwidth, ‘555 channel’ with 605-nm peak excitation/26-nm half-bandwidth, and ‘647 channel’ 706.5-nm peak excitation/36-nm half-bandwidth. For each well, 9 frames with 11 z-stacks per filter were acquired. Z-stack images were deconvolved using maximum projection and each cycle of images were registered using the ASHLAR software^58^. The images were then visualized using the Omero software. For BT lines, cells were plated in 96-well plates (Ibidi) with 10 µg/ml or 20 µg/ml laminin (3 wells/concentration). p-CyCIF was performed as described above with the exceptions that permeabilization was conducted using 0.5% triton-X for 20 min at RT, blocking was done with Super block buffer for 30 min at RT, and Hoechst 33342 was incubated alongside antibodies each cycle overnight at 4°C in the dark. See **Table S3** for a list of antibodies used. Images were acquired with the GE IN Cell Analyzer 6000 using a 20X/0.75 NA objective lens using the filter set described above. 25 frames were collected per well.

### Expansion Microscopy

Tissues were prepared using previously described expansion microscopy protocols^31^. In brief, after dewaxing 10µm thickness FFPE sections were stained with primary and secondary antibodies and hoechst per standard protocols. Tissues were washed 3 times for 10 min at RT, and then covered with 0.03mg/mL Acryloyl-X, SE (Life Technologies, A20770) suspended in 1x PBS for 3 hrs at room temperature. This solution was removed and tissues were covered in cold gelling solution for 30 min at 4°C. Tissue was then enclosed in a gelling chamber constructed using cover glass (24×60mm, No. 1) and incubated at 37°C in a humified environment for 2 hrs. The top coverslip was removed, and tissue was treated with digestion buffer for 3 hrs at 60°C. Samples were washed with 1x PBS buffer for 10 min at RT, stained with Hoechst, and then washed with 1x PBS for 10 min at RT. For expansion, tissues were washed with excess ddH2O (10x gel volume) 3 times for 10 min. Tissues were then imaged using an IN-Cell Analyzer 6000 instrument using a 60x objective lens.

### 3D confocal imaging

20µm FFPE tissue sections were imaged on a LSM980 confocal microscope (Zeiss) with a 63x/1.3NA objective lens. Fluorescent antibodies were excited using the 405nm, 488nm, 561nm, 633nm, and 750nm laser lines. The detection of each channel was separated into individual tracks and detectors to minimize spectral overlap but with Frame Fast to increase acquisition speed. Imaging was confined to selected regions of interest by using Tile Scan with the recommended number of focus support points to approximate a plane. Z stacks were acquired for each tile with a step size of 0.150 µm which was optimized for a pinhole of 1AU. Pixel resolution in X and Y were 0.035 µm. Bit depth was 16 bit and an averaging of 1 was used. Files were saved as czi files and only the first cycle was stitched in ZEN Blue. For subsequent cycles, the raw tiles were registered to this stitched first cycle similar to Nirmal, Vallius, Maliga et al. 2022^26^. Finally, the stitched and registered data was saved as TIFF file and converted to .ims for visualization in Imaris 10.0 (Bitplane).

### Super-resolution imaging

20µm FFPE tissue sections were imaged on a LSM980 confocal microscope (Zeiss) with a 63x/1.3NA objective lens. Fluorescent antibodies were excited using the 405nm, 488nm, 561nm, 633nm, and 750nm laser lines. The detection of each channel was separated into individual tracks and detectors to minimize spectral overlap. Imaging was confined to selected regions of interest by using Tile Scan with the recommended number of focus support points to approximate a plane.

### Focused Ion Beam Serial Electron Microscopy (FIB-SEM)

Tissue samples were fixed with PIPES buffer, stained with osmium tetroxide, and embedded in resin as described in Sheu et al., 2022^59^. Samples were then mounted on aluminum stubs, and the surface was polished using an ultramicrotome and coated with a 20 nm layer of carbon. Images were acquired using the FIB-4 FEI Helios 660 with a 7 nm pixel size and 12 µm milling depth with 30 kV beam and 2.5 nA current. A 35×27×12µm volume was imaged every 20 nm, resulting in 1309 images.

## QUANTIFICATION AND STATISTICAL ANALYSIS

### Image processing and data quantification

Image analysis was performed with the Docker-based NextFlow pipeline MCMICRO 31 and with customized scripts in Python, ImageJ and MATLAB. All code is available in GitHub (https://github.com/labsyspharm/CRC_atlas_2022). Briefly, after raw images were acquired, stitching and registration of the different tiles and cycles was performed with MCMICRO using the ASHLAR module. The assembled OME.TIFF files from each slide were then passed through quantification modules. For background subtraction, a rolling ball algorithm with 50-pixel radius was applied using ImageJ/Fiji. For segmentation and quantification, UNMICST2 was used supplemented by customized ImageJ scripts to generate single-cell data. More details and source code can be found at www.cycif.org and as listed in the software availability section.

### Statistical analysis

Statistical tests are performed using MATLAB (R2023b) and R (R Foundation for Statistical Computing 4.2.2). Two-tailed, two-sample t-tests are performed to compare protein expression between populations within the data from The Cancer Genome Atlas (TCGA) Pan-Cancer Study and TMA cores from IDH-WT GBM. Statistical significance is established when p < 0.05. Pearson correlation coefficient is calculated to determine the linear dependence and p-value is calculated to test the null hypothesis of no relationship between nuclear rupture rate and core level Lamin expression.

### Single-cell data quality control for CyCIF

CyLinter^60^ was used to identify and remove cells affected by visual and image-processing artifacts in single-cell data tables produced by the MCMICRO image assembly and feature extraction pipeline. Specifically, image regions of interest affected by noise (e.g., tissue bunching, antibody aggregates, edge artifacts, necrotic areas, and cells lying outside of the focal plane or grossly mis-segmented) were selected and used to redact the noisy single-cell data from the raw data tables. CyLinter’s cycle correlation filter was then applied to remove cells that shifted or became detached from the microscopy slide over the course of multi-cycle imaging. Post-QC data tables were used in all downstream analyses.

### BAF segmentation

MATLAB was used to segment BAF Foci. Standard deviation filtering of the raw fluorescence intensity was used to capture BAF foci which was normalized against the average core intensity. A binary mask of foci was generated through a constant threshold of 0.8 and assigned to each segmented nucleus. Multiple foci were permitted on the same nucleus.

### Lineage assignment

SOX2-positive GBM cells from post-QC glioma TMA and WSI data were projected onto the four quadrants of a 2D plane representing OPC-, NPC-, MES-, and AC-like states according to the the following GBM cell lineage markers: MAG and SOX10 (OPC-like); ELAVL4HuD and DCX (NPC-like); SOX9, GFAP, EGFR, HOPX, and AQP4 (AC-like); CEBPB, NDRG1, HIF1a, and HIF2a (MES-like). Marker intensities were rescaled (0-1) using the StandardScaler function in Scikit-Learn after removal of upper (99^th^) and lower (1^st^) percentile outliers such that the level of each marker among remaining cells z fell within the range 0z1. Four (4) cell lineage scores were calculated on a per cell basis corresponding to the aforementioned four canonical GBM cell lineages. To do so, the average z values for markers corresponding to a given cell lineage were first identified. For example, in the case of the OPC-like state, raw lineage scores were computed as such: XOPC=12zMAG+zSOX10 .

Thus, values XOPC, XNPC, XAC and XMES were obtained for each cell. Finally, a specific cell lineage score was computed by subtracting the average of four raw lineage scores from specific raw lineage scores of the i^th^ cell. For example, SOPC=XOPC-X in the case of the OPC-like lineage, where X represented the mean across four lineage scores X =14XOPC+XNPC+XAC+XMES. Butterfly plots were generated by projecting final cell lineage scores onto the four quadrants of the 2D plane whose x, and y axes were represented by DX, DY, respectively, and calculated as follows. First, DY was calculated using DY=maxSOPC, SNPC-maxSAC, SMES. DX was then given by SNPC-SOPC when DY>0 and SMES-SAC when DY<0. Thus, the four quadrants I, II, III, and IV represented NPC-, OPC-, AC-, and MES-like GBM cell lineages, respectively.

### BAF Intensity tracking of Live-Cell Imaging Data

Similar to BAF segmentation in CyCIF samples, rupture foci in Kuromochi live-cell time-lapse data was segmented frame by frame. A region of interest is manually selected for each rupture event. The mean nuclear intensity within the ruptured foci was tracked starting from the beginning of the rupture events until the BAF foci resolved. Foci that are not able to resolve within the 20 hrs timeframe or rupture events that happened prior to the start of the capture are disregarded in this analysis. Rupture events in micronucleus, which can be clearly distinguished from the primary nucleus with its smaller size and mobility, are counted separately from the primary nuclear ruptures.

### Focused Ion Beam Serial Electron Microscopy Analysis

FIB-SEM images were cropped, color inverted, despeckled, and registered using ImageJ. Manual segmentation of nuclei was performed for 884 images across 17.7 µm depth in z-direction. Areas are classified into the nucleus, ruptured area, and mitochondria around the rupture site. Volume Annotation and Segmentation Tool (Vast) is used for drawing annotation and 3D rendering.

### Single-Cell RNA-Seq Analysis

Single-cell RNA-sequencing analysis of 28 adult and pediatric high-grade gliomas were performed using data from Neftel et al., 2019^46^. Lineage plots were generated using the Broad Institute Single Cell Portal interface (https://singlecell.broadinstitute.org/single_cell/study/SCP393/single-cell-rna-seq-of-adult-and pediatric-glioblastoma#study-visualize). The processed log TPM gene expression matrix, metadata file, and the hierarchy data were downloaded from the Broad Single Cell portal (https://singlecell.broadinstitute.org/single_cell). Further analyses were performed in R (v4.2.2) using the Seurat (v4.2.0), ggplot2 (v3.4.2), ComplexHeatmap (v2.14.0), and ggsignif (v0.6.4) packages.

### Blinding

For quantification of BAF foci in human tissues in human tissue microarrays and cell lines, the reviewing pathologist (S.C.) and analyses were blinded to tumor subtype and patient data. Analysis of CyCIF data, including lamin, cGAS, pH2AX, and BAF quantification was blinded to tumor subtype and patient data.

