## Supplementary figures and images for "2D and 3D multiplexed subcellular profiling of nuclear instability in human cancer"

### Supplemental Figures

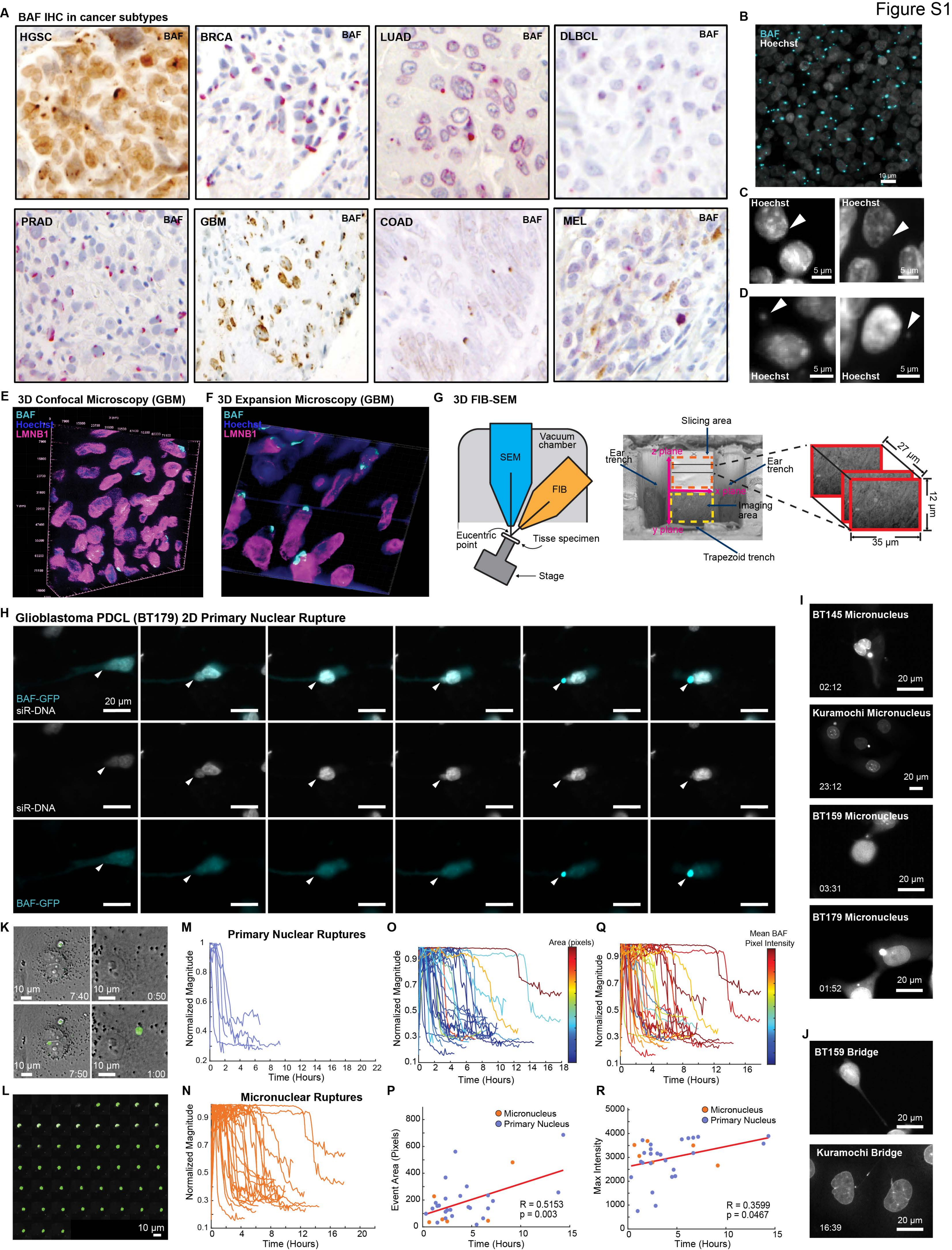

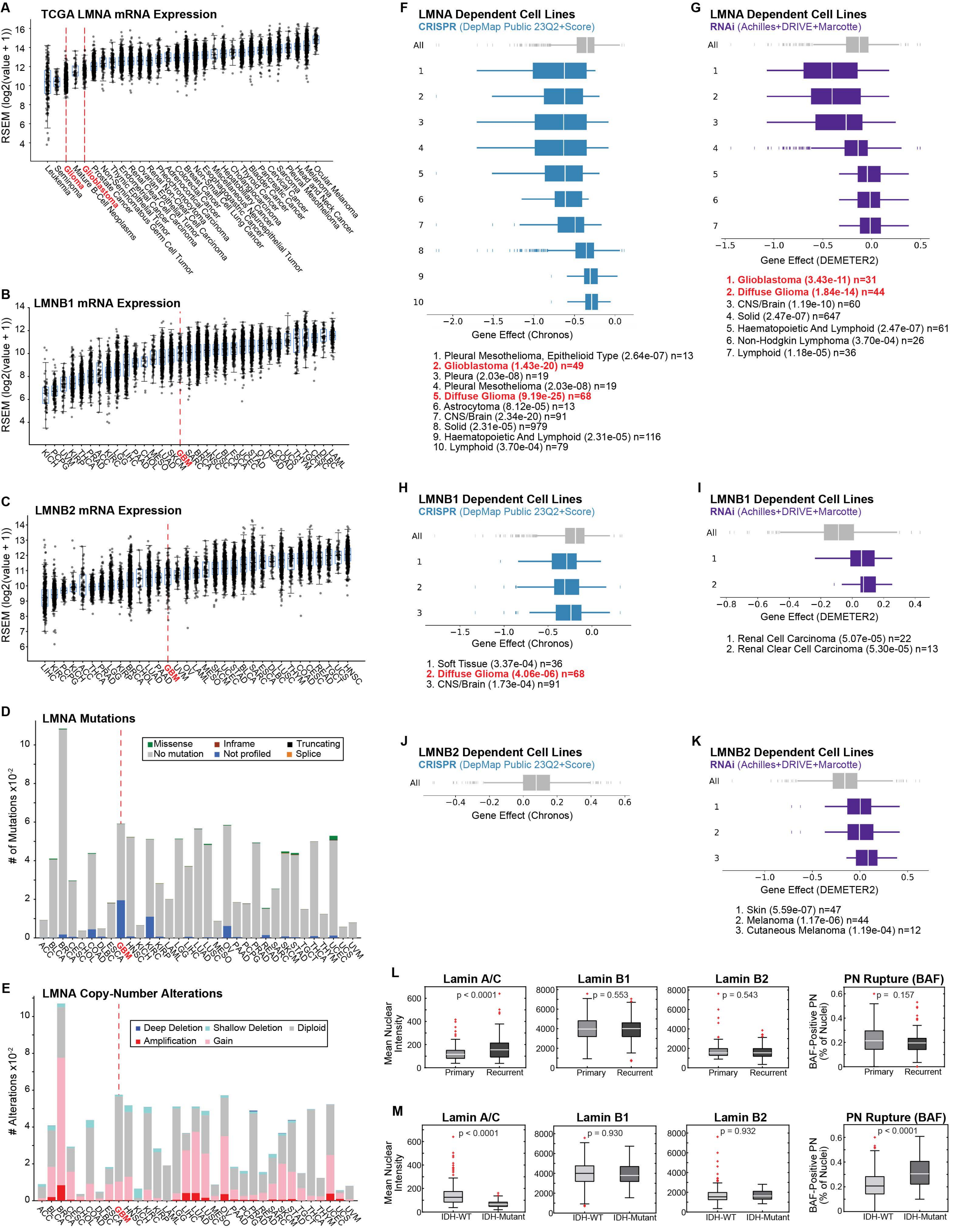

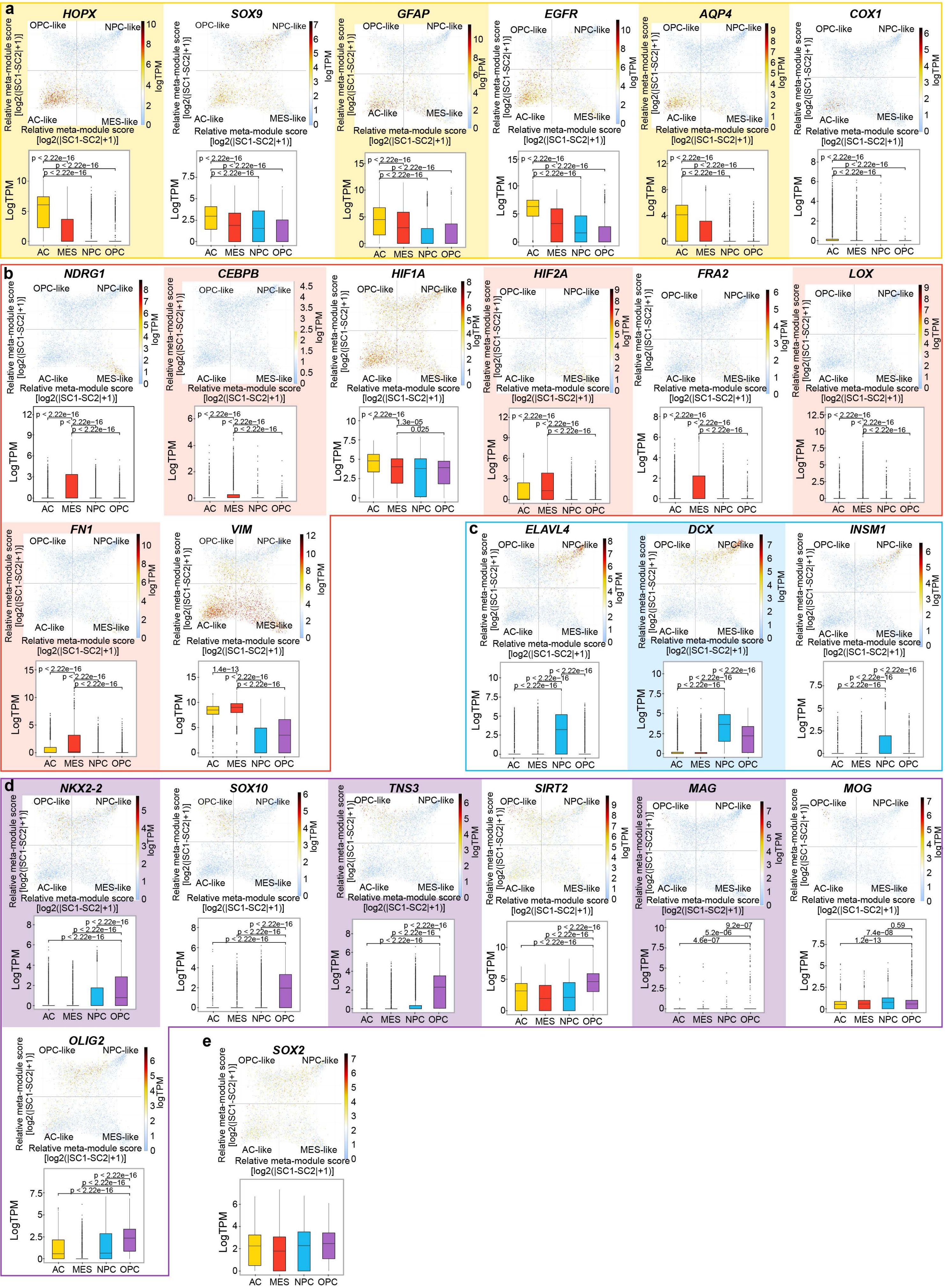

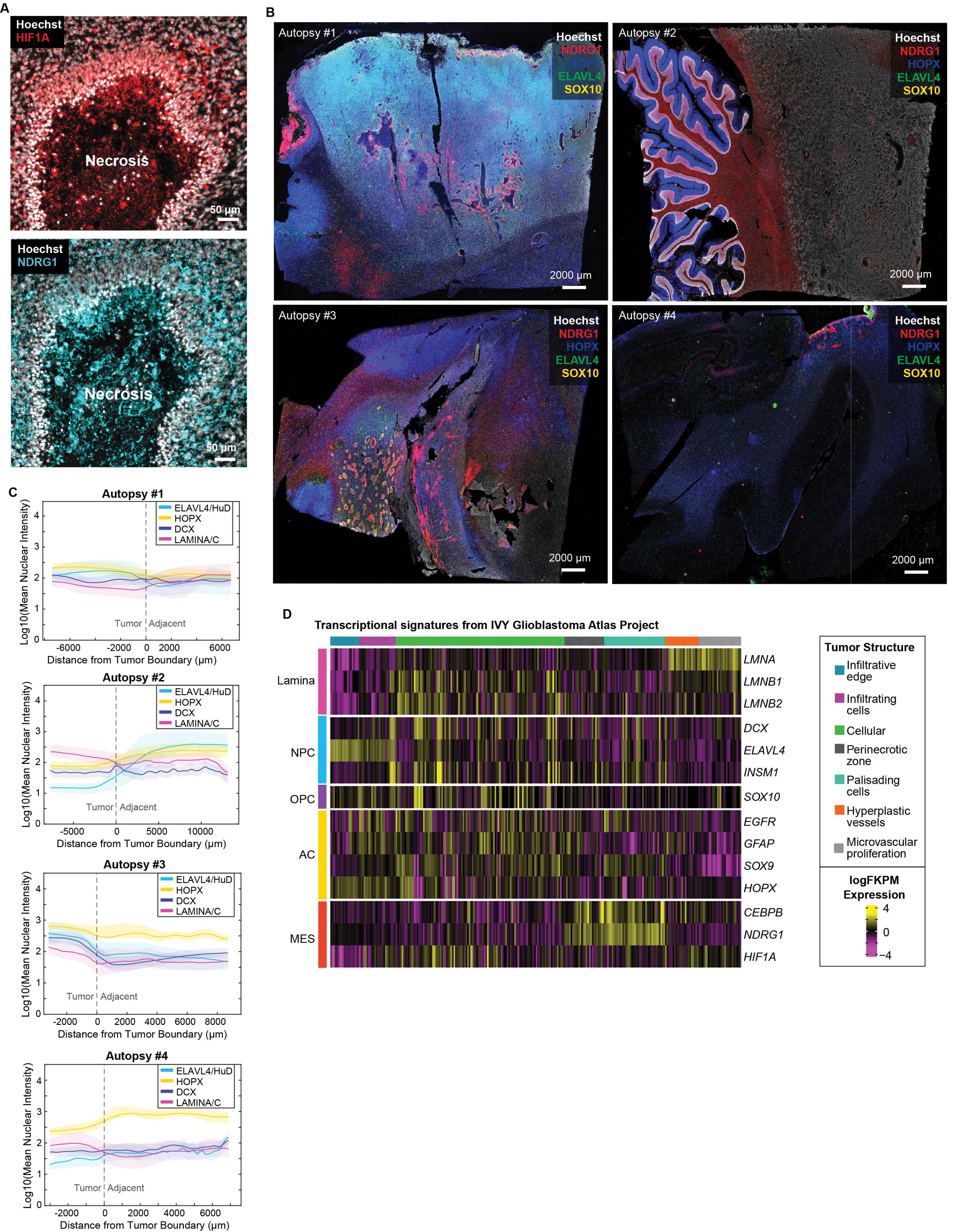

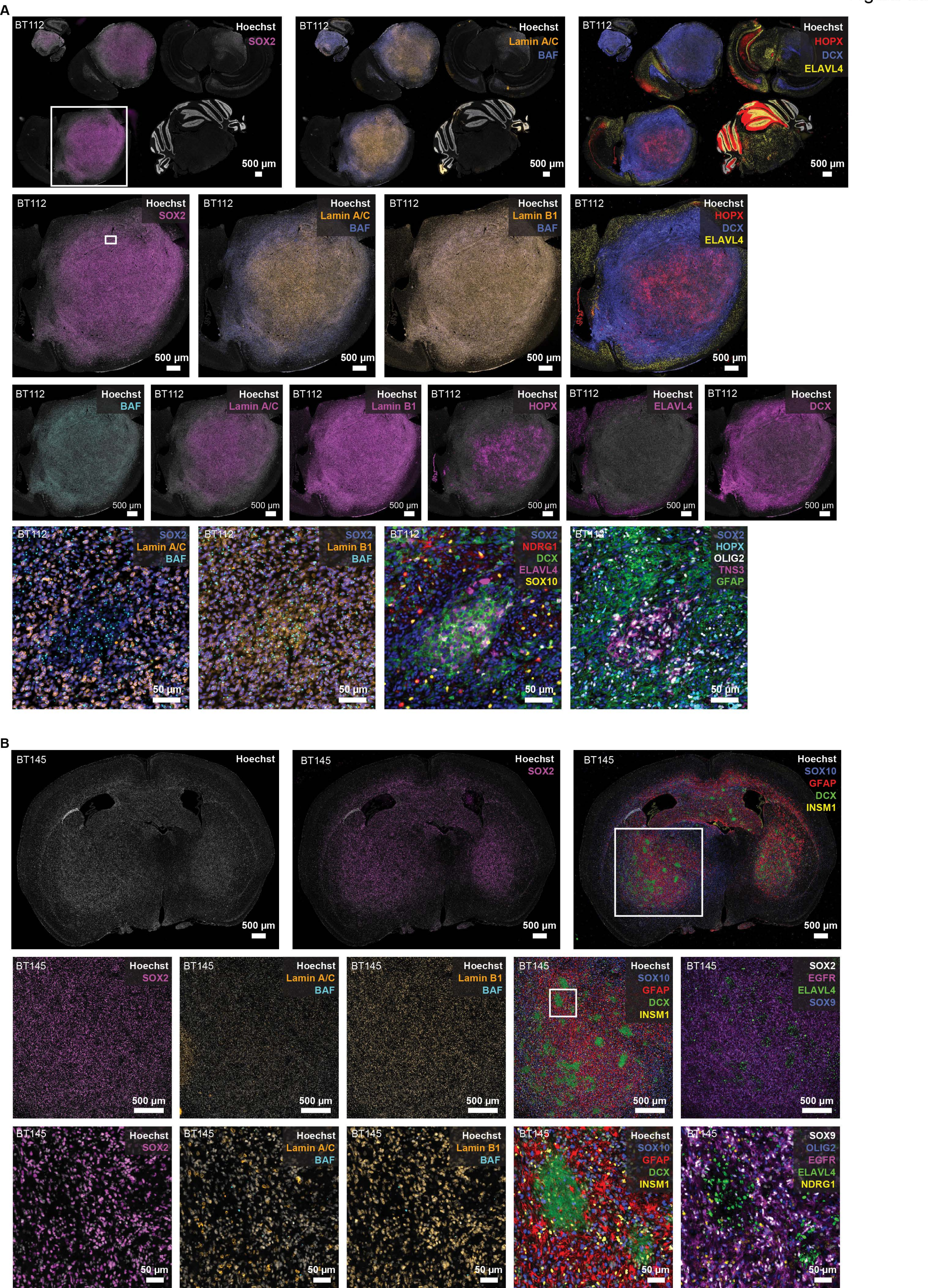

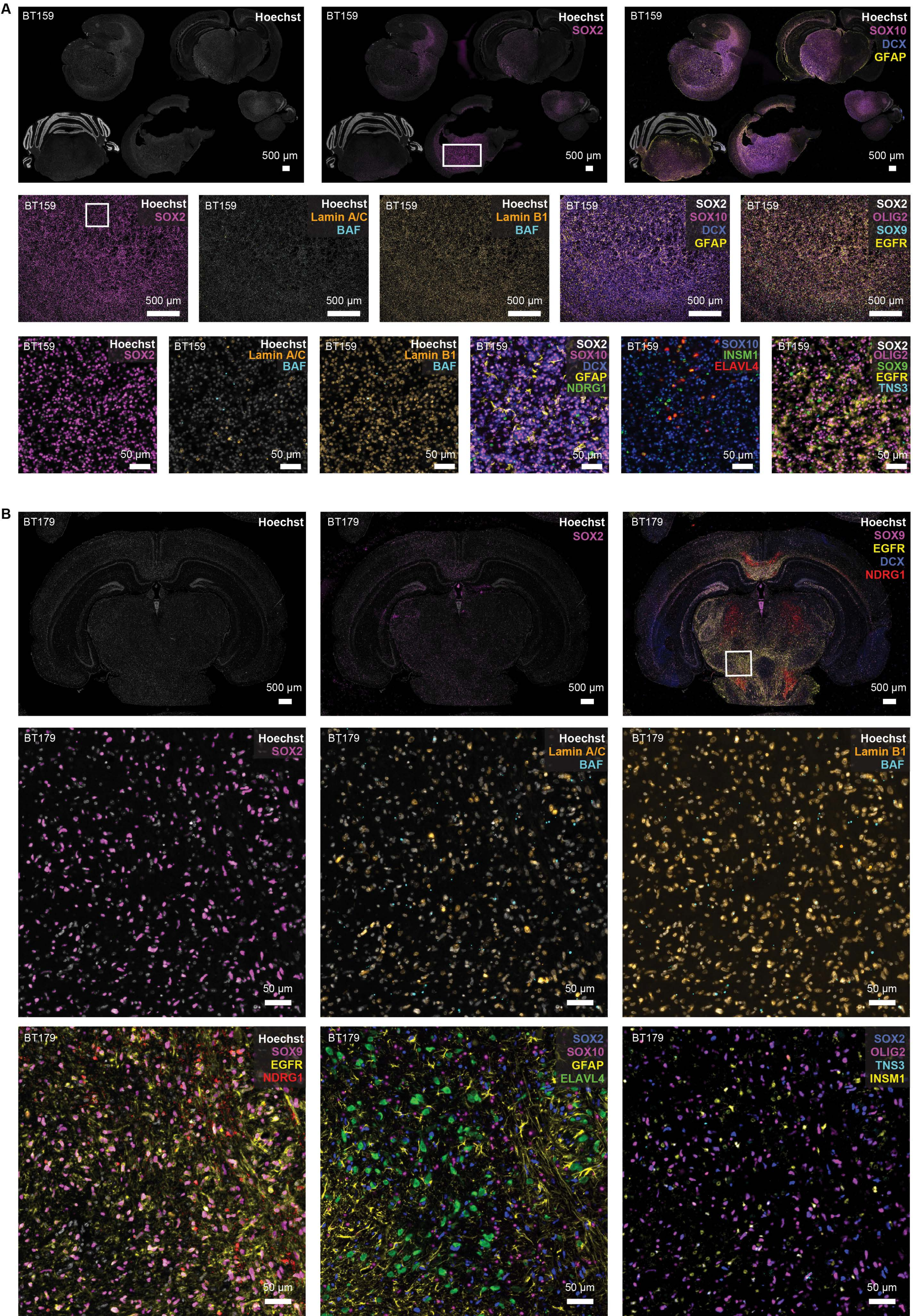

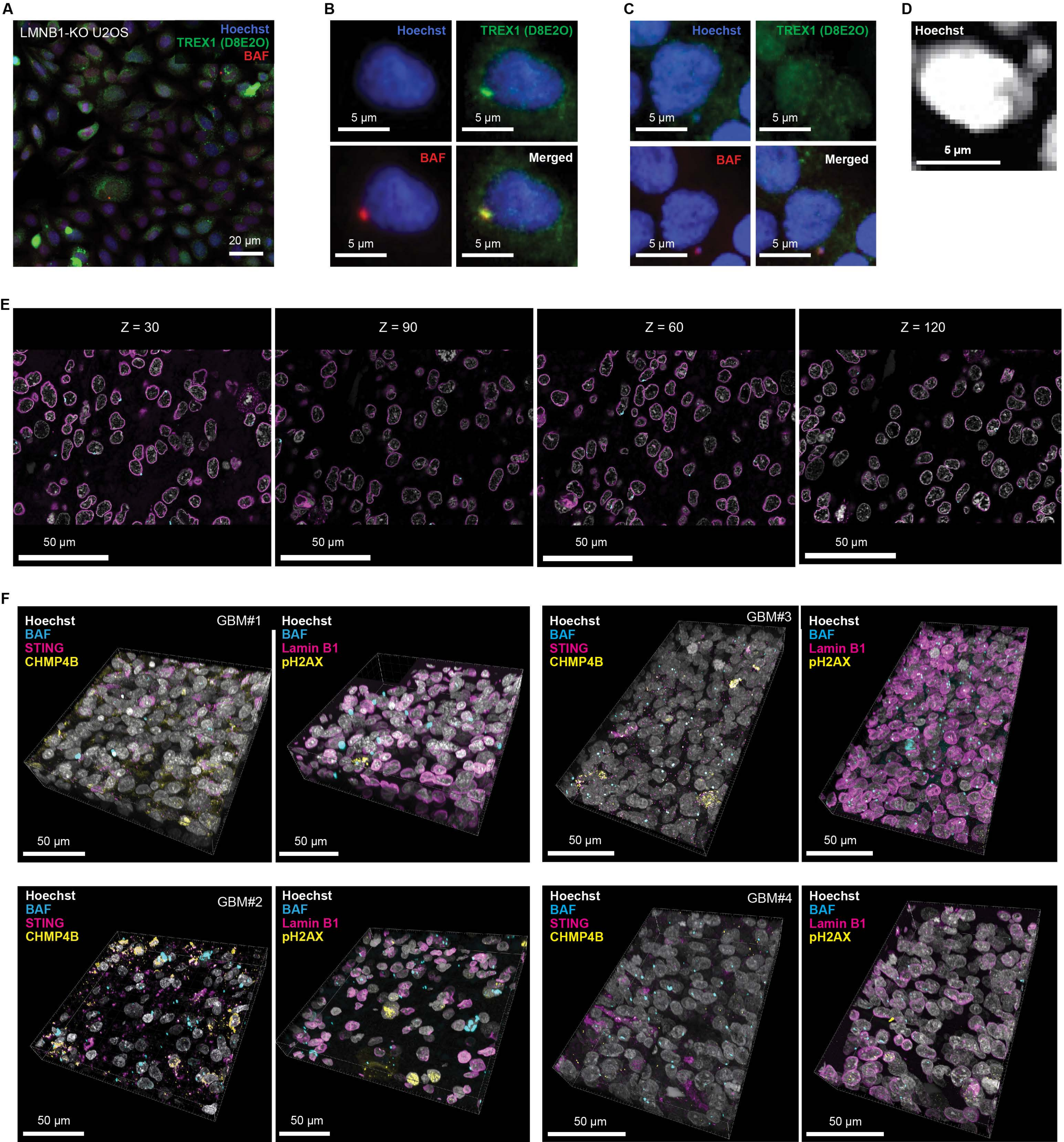
